# DNA methylation and gene expression trajectories of human postprandial metabolism

**DOI:** 10.1101/2024.11.29.626085

**Authors:** Ricardo Costeira, Lidia Daimiel Ruiz, Thies Gehrmann, Fatih Bogaards, Sergio Villicaña, Lucy Sinke, Yasrab Raza, Max Tomlinson, Colette Christiansen, Bastiaan T Heijmans, P Eline Slagboom, Tim D Spector, Kerrin S Small, Juan F Alcala-Diaz, Oriol Rangel-Zuñiga, Jose López-Miranda, Melanie Waldenberger, Sarah E Berry, José M Ordovás, Jordana T Bell

## Abstract

Human postprandial metabolism is characterised by a highly individualised response to food that is predictive of cardiometabolic health and underexplored at the molecular level. We profiled blood DNA methylation (DNAm) and gene expression trajectories before and after a test meal in 225 European participants. We identify DNAm changes at fasting, 30 minutes and 4 hours after meal challenge, including in metabolically relevant genes *INPP4A, GHRL, ASIP* and *ABCG1*, with changes observed as early as 30mins postprandially. Gene expression trajectories also changed postprandially predominantly at 4 hours, with replication of lipid metabolism (*CPT1A)* and circadian rhythm (*PER1)* genes. Genetic variants affect postprandial molecular trajectories at genes linked to obesity (*PDE9A)* and glucose response (*GPT2)*. Multiple signals associated with postprandial glucose and triglyceride levels, with replication of *CPT1A* methylation. The postprandial DNAm and expression trajectories target metabolically relevant genes, giving insights towards mechanisms underlying inter-individual response to food and cardiometabolic disease risk.

## Introduction

The postprandial state, defined as the 0–8-hour period following a meal, is characterised by highly individualised responses to food in humans^1^. Intra- and inter-individual variation observed in the glycemic and lipemic responses to food are associated with poor cardiometabolic health outcomes, including type 2 diabetes and an increased risk of cardiovascular events^2^. Postprandial oxidative stress and inflammation partly mediate the acute effect of foods on disease risk^2^. However, the molecular mechanisms associated with the postprandial state in humans are still not well understood. Epigenetic regulation of gene expression, particularly through DNA methylation (DNAm) at CpG sites, may play a role in postprandial metabolism.

DNAm is a major regulator of gene expression in humans^3^, and is responsive to various stimuli, including food^4^. Despite this, human studies on postprandial DNAm are limited. So far, one larger scale study identified DNAm signals at fasting to be associated with postprandial triglyceride concentrations^5^, highlighting key metabolic genes such as *CPT1A, ABCG1,* and *SREBF1.* Two smaller scale studies further explored DNAm longitudinally over the postprandial phase, in 26 healthy males at baseline fasting and 160 minutes (m) postprandially^6^, and in 12 males across different BMI categories from baseline up to 10 hours (h) postprandially^7^. The results revealed that changes in blood DNAm are partially mediated by shifts in blood cell proportions and that DNAm associations with BMI and fatty acid concentrations differ between fasting and postprandial states.

While DNAm studies offer insights into the epigenetic landscape of the postprandial state, transcriptomic research has also identified changes in postprandial gene expression. Multiple studies have identified changes in postprandial expression levels of immune, inflammatory and circadian rhythm genes, such as *PER1,* and body-weight dependent metabolic responses to food, but results are often context-specific and limited to small samples^8–12^. More recently, the larger Growing Old Together study (GOTO)^13^ identified postprandial blood gene expression levels from fasting to 30m postprandially ^14^. This was carried out both pre- and post- a combined nutritional and physical activity lifestyle intervention in 85 participants, and identified changes in stress response genes, with stronger effects observed in males ^14^.

Despite significant advances, a comprehensive understanding of how DNAm and gene expression change over the postprandial state remains elusive, particularly in large and diverse human populations. In this study, we characterize human blood DNAm and gene expression trajectories at fasting and over the postprandial state (30m to capture peak glycemia, and 4h for peak lipemia) in 225 participants from two large-scale European postprandial cohort studies, PREDICT and CORDIOPREV. We identified over a hundred molecular genomic signals that changed after test meal and characterized their links to postprandial glucose and triglyceride trajectories, which are cardiometabolic risk factors.

## Results

We captured longitudinal DNAm and gene expression signals at fasting and postprandially at 30m and 4h after test meal (Figure 1). DNAm differences between fasting baseline and 4h postprandially were explored by meta-analysis of the PREDICT and CORDIOPREV studies in up to 225 participants (54% males, median age = 58.8 years). The PREDICT study is a large-scale high-resolution postprandial study with 1002 participants recruited at baseline from the UK population^15^, where healthy participants were fed a high carbohydrate-high fat meal. We generated whole blood DNAm profiles in a subset of individuals from the PREDICT study, including 104 PREDICT female healthy participants (mean age = 58.97 ± 12.58 years), at fasting and postprandially at 30m and 4h after meal intake. CORDIOPREV (the CORonary Diet Intervention with Olive Oil and Cardiovascular PREVention study) is a randomized clinical trial involving 1002 patients with coronary disease^16^, where participants received a high-fat meal at baseline to assess postprandial lipemic response. We generated whole blood DNAm in a subset of CORDIOPREV individuals, including 121 male participants (mean age = 59.09 ± 9.48 years) at fasting and 4h after meal intake (Table 1). DNAm trajectories were further dissected in the PREDICT cohort alone, including 30m postprandial glycemia. Matched RNA-sequencing expression trajectories were generated in a random subset of 50 PREDICT females to explore postprandial gene expression changes. DNAm and expression trajectories were then linked to the trajectories of plasma glucose and blood triglyceride concentrations after test meal in PREDICT. Replication of the postprandial regulatory and functional genomic trajectories was undertaken using epigenetic results from postprandial studies by Lai et al.^5^ and Rask-Andersen et al.^6^, as well as gene expression postprandial results from the lifestyle intervention study by Gehrmann et al^14^.

**Figure 1.**
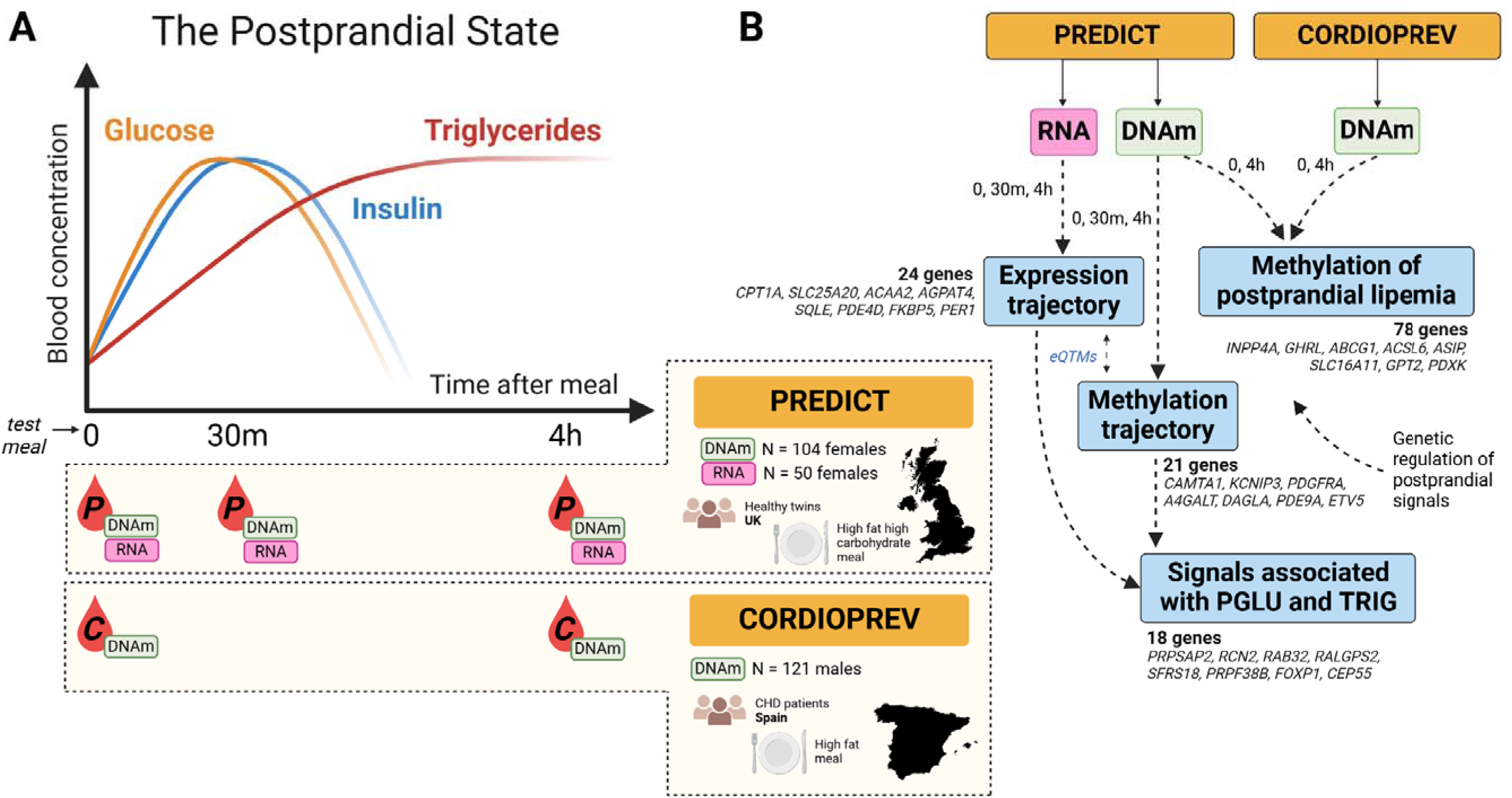
Sampling strategy over the postprandial state (**A**) and the main analyses and results from this study (**B**). Analyses were performed using blood samples from the PREDICT and CORDIOPREV cohorts at fasting baseline, 30m and 4h after meal.

**Table 1.** Study sample characteristics (n = 225).

|  | PREDICT | CORDIOPREV |
| --- | --- | --- |
| n of participants | 104 females | 121 males |
| n of DNAm samples (fasting, 30m, 4h) | 102, 103, 103 | 117, 0, 115 |
| n of gene expression samples (fasting, 30m, 4h) | 50, 49, 49 | – |
| Age (mean ± sd) | 58.97 ± 12.58 | 59.09 ± 9.48 |
| BMI (mean ± sd) [kg/m<sup>2</sup>] | 25.64 ± 5.28 | 30.95 ± 4.24 |
| Smoking (never, ex, current) | 58, 42, 4 | 24, 81, 16 |

### Longitudinal meta-analysis identifies DNA methylation signals postprandially at 4 hours

Postprandial lipemia is an independent risk factor for cardiovascular disease ^17^. It is characterized by a rise in blood lipids after meal consumption, which peaks around 4h and returns to baseline around 8h postprandially. We aimed to identify DNAm changes that occur at this timepoint and are potentially characteristic of postprandial lipemia. To this end, we contrasted fasting baseline and 4h post-meal DNAm levels in the PREDICT and CORDIOPREV cohorts. Meta-analysis of epigenetic changes between 4h and baseline identified 108 DMPs in 78 genes at FDR 5% and after heterogeneity filters (HetISq < 85% and HetPval ≥ 0.01), across the two cohorts (Figure 2A, Figure S1 and Table S1). The peak signal was observed at cg03403093 in the 5’UTR region of *INPP4A* (coefficient *b* = 0.121 ± 0.020, *p* = 1.98E-09), a lipid phosphatase that negatively regulates the PI3K/Akt pathway^18^. Other significant CpG signals were found in metabolically relevant genes such as *ACSL6, GHRL, ASIP, SLC16A11, PDXK, ABCG1, GPT2* and others (Figure 2 and Figures S2, S3). These include lipid metabolism genes, such as *ASIP* and *ABCG1*, a key lipid transporter^19^, as well as the acyl-CoA synthetase *ACLS6* and *SLC16A11* transporter, implicated in lipid synthesis and metabolism^20,21^. *GHRL* encodes the prepropeptide for ghrelin, an appetite-stimulating hormone with effects in growth hormone release from the anterior pituitary, adiposity, circadian regulation and metabolic inflammation (metaflammation)^22,23^. Altogether, the *ASIP*, *ABCG1, GHRL, ACSL6, SLC16A11, GPT2* and *PDXK* have all been studied in the context of obesity and type 2 diabetes^19–26^, while the PI3K/Akt pathway is the main signalling pathway implicated in the relationship between these two traits^27^. Biological pathway and enrichment analyses of our results highlighted pathways for neuronal system activity, signal transduction and fatty acid metabolism (Table S2). Enrichment analysis of phenotypes previously linked to these CpGs in the EWAS Catalog^28^ identified enrichment for signals previously associated with BMI, waist circumference, C-reactive protein levels, and granzymes A and K (GZMA and GZMK) levels (Table S3). C-reactive protein is a marker of inflammation, while GZMA and GZMK are cell death–inducing enzymes stored in cytotoxic T lymphocytes and NK cells^21^. Other metabolic phenotypes were also linked to these CpGs in previous EWAS within the EWAS Catalog, including type 2 diabetes, and levels of glucose, triglycerides and cholesterol, but these traits did not show evidence for enrichment.

**Figure 2.**
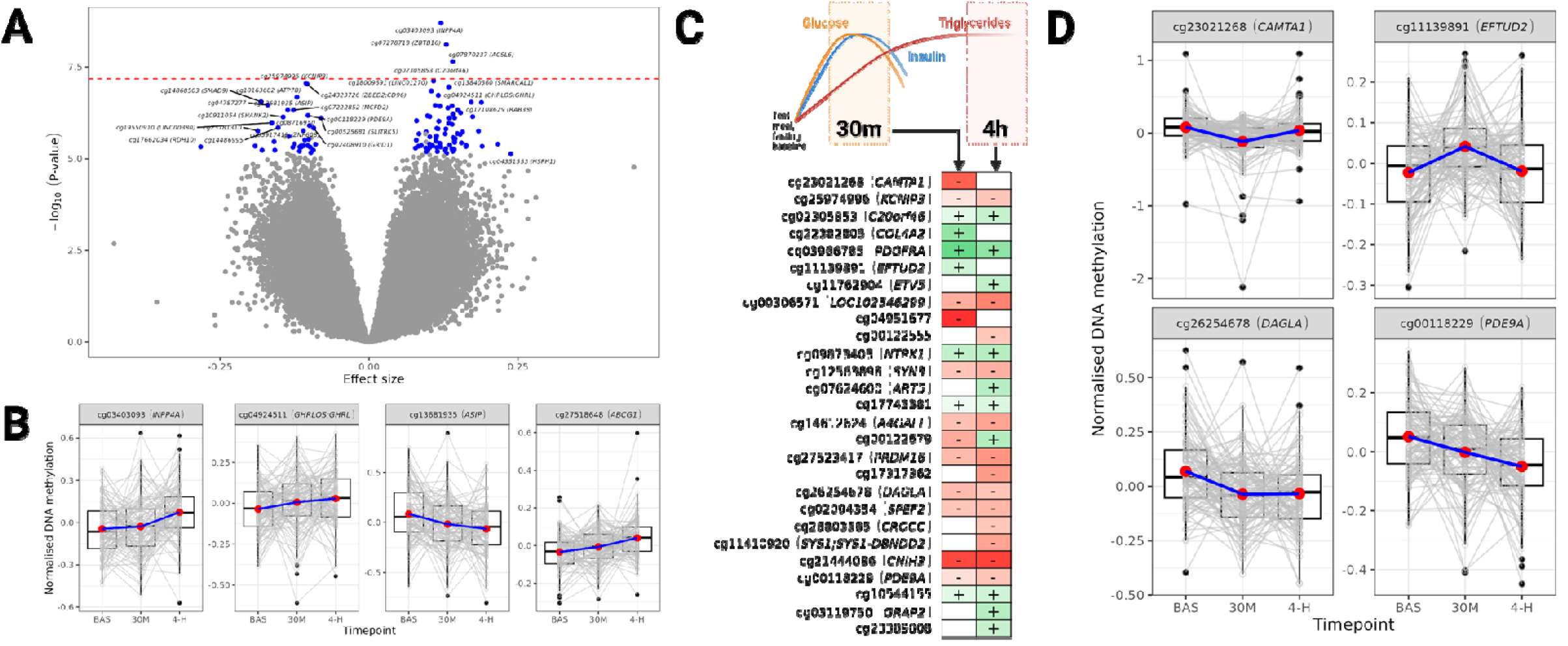
DNAm signals identified postprandially. **A.** Signals associated with postprandial lipemia at 4h in PREDICT and CORDIOPREV (n = 225 participants). Signals in blue passed the methylation meta-analysis heterogeneity filters and multiple testing correction threshold FDR = 5%. Signals above the dashed red Line passed the epigenome-wide Bonferroni P-value correction threshold based on 746,953 autosomal probes. The effect sizes displayed are based on DNAm M-values and the signals represent the sex-shared effects observed across PREDICT and CORDIOPREV. **B.** Trajectory of peak signals identified from the 4h meta-analysis in the PREDICT sample. **C.** Signals identified in the PREDICT sample across fasting baseline, 30m and 4h postprandially (n = 104 participants). Signals are ordered by their significance P-value and heatmap intensity corresponds to strength of effects. Positive and negative directions of effect are marked green and red, respectively, while non-significant effects were left blank in the heatmap. DNAm trajectories are reported with FDR = 5%. **D.** Examples of the shape of different DNAm trajectories identified from longitudinal DNAm analysis in PREDICT.

Like previous work (Rask-Andersen et al.^6^) we observed postprandial changes in blood cell composition in our samples (Figure S4). Specifically, the proportions of monocytes, granulocytes and natural killer cells changed significantly after test meal. In the epigenome-wide cohort-specific analyses, and meta-analysis, results are presented after adjustment for blood cell proportion changes. Furthermore, we profiled DNAm in CD4+ T cells from a random subset of 10 PREDICT participants (6 females and 4 males, mean age = 75.91 ± 6.12 years, mean BMI = 24.23 ± 1.36 kg/m^2^) and evaluated the consistency of effects across whole blood and CD4+ T cells postprandially. Of the signals that show the same direction of effect in whole blood at 4h across PREDICT, CORDIOPREV, and the subset of 10 PREDICT participants, 11 signals (41%) exhibit the same direction of effect across whole blood and CD4+ T cell populations (Table S4).

As a validation of the epigenome-wide meta-analysis results we also carried out analyses using unadjusted DNAm β-values. These altogether identified 175 DMPs, including 70% of signals from the main analysis based on M-values (113 DMPs total, FDR = 5%), but showed greater evidence for heterogeneity in meta-analysis. Directions of effect were overall consistent between M- and β-values (Figure S5). Further validation analyses in the complete sample of 265 individuals included analyses of the complete set of female and male participants across both cohorts (Methods), and this identified 20 DMPs (FDR = 5%), 19 of which are in the primary meta-analysis results presented above.

### Dissection of DNA methylation and expression trajectories over the postprandial state

The CORDIOPREV study captured the response to dietary fat in coronary heart disease patients from baseline fasting to 4h postprandially alone ^16^. In contrast, the PREDICT study captured the human response to food^1^ from baseline fasting to both the glycemic response at 30m and the lipemic response at 4h. We therefore explored DNAm and gene expression across fasting baseline, and postprandially at 30m and 4h in the PREDICT sample alone, in order to explore the full DNAm and expression trajectories after meal.

#### Postprandial DNA methylation trajectories

First, using the 3 timepoints from PREDICT we explored how early after test meal the 108 DMPs captured earlier at 4h (Table S1) show differential DNAm in comparison to fasting DNAm levels. Of the 108 DMPs captured from PREDICT and CORDIOPREV at 4h, 95% were captured at 4h in the PREDICT trajectory analysis alone, and a subset of 40% DMPs showed DNAm changes as early as 30m after test meal (*p* < 0.05; Table S5). Signals with nominal changes at 30m included cg07870237 (*ACSL6*), cg13681935 (*ASIP*) and cg14880584 (*GPT2*). To further dissect DNAm changes 30m after meal we extended our full trajectory analysis in PREDICT epigenome-wide.

Variability in DNAm levels across 3 timepoints (fasting baseline, 30m and 4h postprandially) was identified at 27 CpGs epigenome-wide, which annotated to 21 genes including *CAMTA1, KCNIP3, PDGFRA, A4GALT, DAGLA, PDE9A* and *ETV5* (Figure 2C and Table S6: FDR = 5%). The peak signal was cg23021268 in the body of *CAMTA1* (*b* = −0.314 ± 0.053, *p* = 5.10E-08), which encodes a transcription activator that regulates insulin production and secretion^29^. DAGLA is a diacylglycerol lipase, *A4GALT* is upregulated with high salt diet ^30^, and *ETV5* and *PDE9A* expression is associated with obesity in mice ^31,32^. For the majority of signals DNAm levels changed immediately 30m after meal intake, with variable patterns over the postprandial state (Figure 2D). Notably, *CAMTA1* DNAm was lowest at 30m highlighting its potential role in the glycemic response. *DAGLA* gene hypomethylation at cg26254678 was sustained up until 4h after a meal. Altogether, there were 108 DMPs in the meta-analysis of PREDICT and CORDIOPREV at 4h, and 27 DMPs in the postprandial DNAm trajectory analysis in PREDICT females alone, of which six signals overlapped. The six overlapping DMPs included cg02305853 (*C20orf46*), cg09873405 (*NTRK1*), cg10544155, cg25974996 (*KCNIP3*), cg00118229 (*PDE9A*) and cg00122255. At gene level, postprandial changes in DNAm signals were detected in 78 genes in the meta-analysis of PREDICT and CORDIOPREV at 4h, and in 21 genes in the postprandial DNAm trajectory analysis in PREDICT females alone, of which four genes overlapped; resulting in 95 unique genes that exhibit postprandial DNAm changes.

We conducted several sensitivity analyses, first, exploring the consistency of DNAm trajectories both at these 27 CpGs and at the 108 signals from the postprandial 4h meta-analysis across different DNAm quantifications. We observed that results, based on DNAm M-values, were overall consistent with the trajectories of unadjusted raw DNAm β-values (Figures S2-S3, S6-S7). Second, trajectories of DNAm were also consistent across whole blood and CD4+ T cells only in a subset of CpGs (Figure S8). Because the PREDICT sample used was composed of predominantly females (Table 1), a sensitivity analysis accounting for menopausal status and hormone replacement therapy was conducted in a subsample of 37 females with available phenotype information. Altogether, twenty-three of the 27 DMPs remained nominally significant and in the same direction of association in this subsample (85% of DMPs, Table S7). Finally, previous research has found that human DNAm levels can oscillate with time of day^33^. To ascertain that the DNAm trajectories observed were due to test meal and not time of day we interrogated the set of circadian DMPs reported by Oh et al.^33^. We were able to interrogate 58% of circadian DMPs and, while a few DMPs were variable at a nominal significance (Table S8), we did not find major evidence for circadian effect on the DMPs identified at 30m and 4h after test meal. No effect of time of day was observed for the 4h postprandial lipemia meta-analysis results either (Table S8).

Gene enrichment analysis of DNAm trajectory results identified 80 terms across 5 ontology/pathway databases, including regulation of transcription, signalling (*e.g.:* PI3K), cellular response to oxidative stress, haematopoiesis, and neurotransmission where 11% of the terms identified were directly related to brain (Table S9). In the EWAS Catalog we observed an enrichment for signals previously linked to immune phenotypes (*e.g.:* C-reactive protein, GZMK, GZMA and IL2RB) and found that some of the CpGs were previously associated with type 2 diabetes, glucose and BMI (Table S10).

We considered replication of the DNAm postprandial findings using previously published results^6^. We initially compared our results to those from Rask-Andersen et al.^6^ in 26 males sampled at baseline and 160m after consumption of a different standardised meal. Twenty-four signals captured at 160m postprandially by Rask-Andersen et al. were nominally significant in the PREDICT sample with 9 (38%) and 13 (48%) of signals matching direction of effect at 30m and 4h in PREDICT (Table S11A), respectively, despite sampling at different time-points postprandially. The peak replicated signal, and only signal to pass Bonferroni correction, was cg08697116 in the body of *KLK4* (*b* = 0.141 ± 0.045, *p* = 4.86e-05) with the same direction of effect at 4h postprandially. The larger meta-analysis of postprandial lipemia DNAm signals replicated 10 other signals from Rask-Andersen et al. at nominal significance (Table S11B), and 4 of the 24 signals already captured by PREDICT alone (Table S11A).

#### Postprandial gene expression trajectories

Variability in longitudinal gene expression trajectories identified 24 differentially expressed genes over the postprandial state in a random subsample of PREDICT individuals (50 females sampled at 0-30m-4h, Figures 3A-B, Figure S9 and Table S12: FDR = 5%). The peak differentially expressed trajectory was observed for *CPT1A*, which was downregulated at 4h (*b* = 0.183 ± 0.119, *p* = 6.60E-25). *CPT1A* encodes the key rate-limiting enzyme of fatty acid oxidation and epigenetic regulation of *CPT1A* expression has previously been observed with high fat^34^. Further differentially expressed signals included the circadian rhythm regulator *PER1,* recently linked to the regulation of metabolic homeostasis in fasting^35^, as well as genes involved in metabolism such as *SLC25A20, ACAA2, AGPAT4* and *SQLE*, and inflammation regulators *PDE4D* and *FKBP5*. While we observed variability in the shape of the postprandial DNAm trajectories (Figure 2D), the gene expression signals were consistently heightened at 4h postprandially. Furthermore, the direction of effect was generally consistent across 30m and 4h (Figure 2, Figure S10 for all trajectories). Gene expression analysis of postprandial lipemia alone in PREDICT (0-4h changes) identified 21 differentially expressed genes, including 18 genes from our main gene expression trajectory analysis (75% of genes from the 0-30m-4h analysis, Table S12).

**Figure 3.**
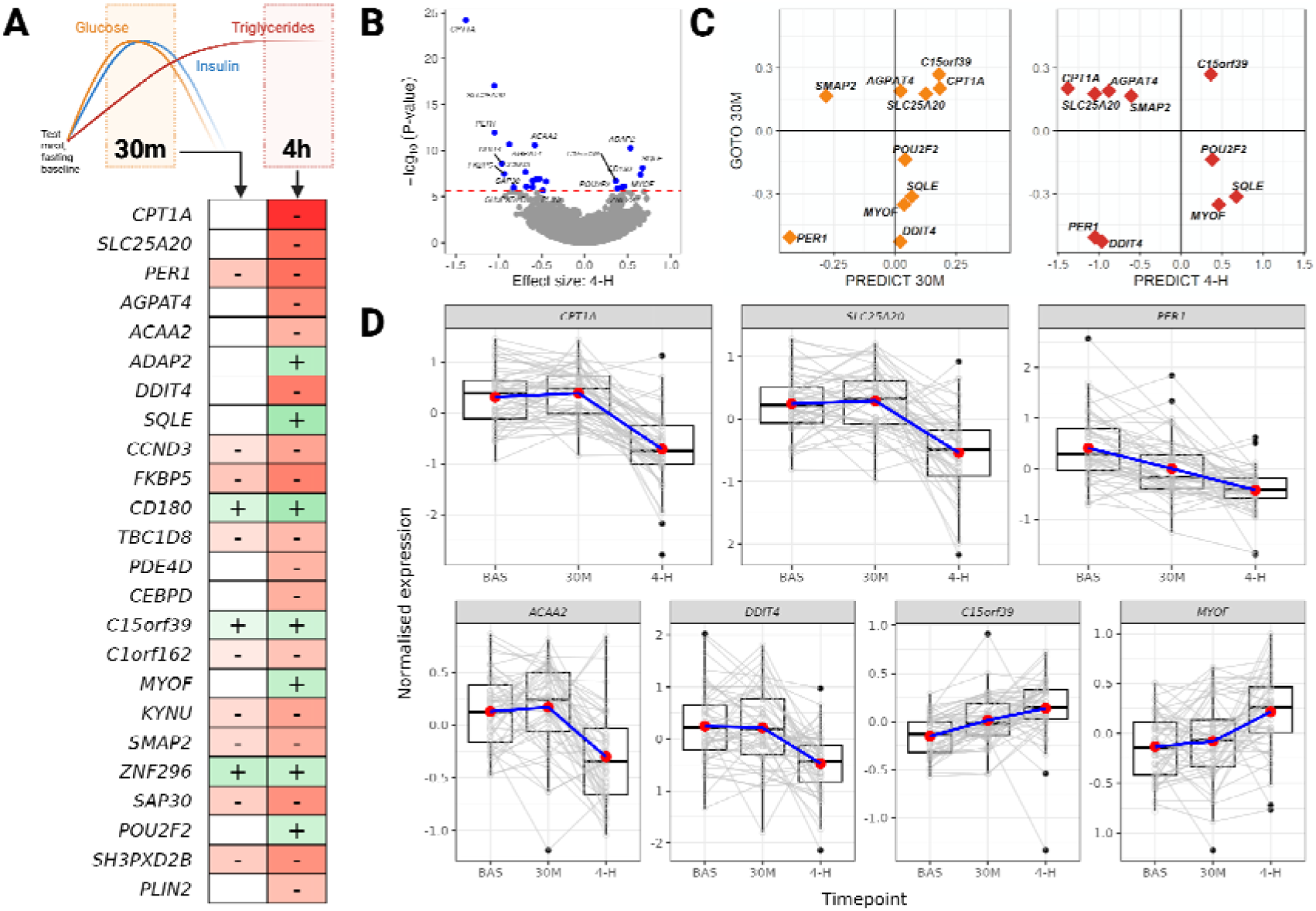
Gene expression signals identified postprandially. **A.** Transcriptome-wide signals identified in the PREDICT sample across fasting baseline, 30m and 4h postprandially (FDR = 5%, n = 50 participants). Expression signals are ordered by their significance P-value and heatmap intensity corresponds to strength of effects. Positive and negative directions of effect are marked green and red, respectively, while non-significant effects were left blank in the heatmap. **B.** Volcano plot of expression signals associated with postprandial lipemia at 4h in PREDICT. **C.** Replication of expression signals in the GOTO cohort. Signals replicated in GOTO at 30m postprandially and their effects in GOTO and PREDICT at 30m and 4h postprandially are represented (adj. *p* < 0.05). **D.** Trajectory of expression signals identified across PREDICT, GOTO and Rask-Andersen et al. (2016) over the 3 timepoints sampled in PREDICT (adj. *p* < 0.05).

Sensitivity analysis was performed for gene expression using the full set of covariates from the epigenetic analyses including estimated blood cell proportions (Methods), and all 24 expression signals remained significant with the same direction of effect (Table S13). Enrichment analysis of the gene expression trajectory genes identified enrichment in cell signalling pathways, insulin response, cholesterol and lipid metabolism, inflammation, haematopoiesis and blood cell differentiation, and others (Table S14).

Replication of the differential gene expression results was pursued in two external datasets. First, we used data from the GOTO lifestyle intervention study in 85 individuals, which includes blood gene expression at the fasting state and 30m after a standardized nutrient challenge, both before and after a combined lifestyle intervention^14^. We replicated 10 out of 24 (42%) PREDICT gene expression signals in the combined male and female sample from GOTO prior to intervention, at Bonferroni correction (adj. *p* < 0.05; Table S15A). Eight genes showed the same direction of effect in GOTO at 30m as in the PREDICT 30m and 4h data postprandially, including *CPT1A* (30m), *SLC25A20* (30m), *AGPAT4* (30m) and *PER1* (30m, 4h) (Figure 3C and Figure S11), which are the peak signals from postprandial gene expression trajectory analyses. Eight of the 10 genes replicated in the GOTO combined analysis were also identified in the GOTO females alone (Table S15B), and 6 were identified post-exercise intervention as well (*PER1, AGPAT4, DDIT4, SQLE, C15orf39* and *SMAP2)*. Other genes with signals replicated post-intervention in GOTO were *ACAA2, FKBP5* and *CCND3* (Table S15).

We also considered whether our results replicate previous blood gene expression findings from Rask-Andersen et al. (2016) at baseline fasting and 160m postprandially, based on microarray profiling. Of the 945 signals tested, we replicate 17 (adj. *p* < 0.05; Table S16). Altogether, *CPT1A, SLC25A20, PER1, ACAA2, DDIT4, C15orf39* and *MYOF* were identified as differentially expressed in blood samples across three independent studies with different test meal challenges – in PREDICT RNA-seq data, in GOTO RNA-seq data from Gehrmann et al^14^, and in the microarray expression data from Rask-Andersen et al.^6^ (Figure 3D).

#### Coordinated DNA methylation and gene expression postprandial signals

We found evidence of coordinated changes in DNAm and gene expression over the postprandial state. In the postprandial trajectory results there were 940 genes for which both DNAm and gene expression changes were observed over the postprandial state at nominal significance (Figure S12A). This corresponds to 52% and 6% of all differentially expressed and methylated genes at nominal significance (*p* < 0.05) postprandially, respectively. Genes that exhibit coordinated DNAm and expression changes included previously reported *ABCG1, DAGLA, ACSL6* and *NTRK1* (Table S17A). Similarly, comparison of 4h meta-analysis DNAm results and gene expression results identified 907 genes that harbour both sets of signals at nominal significance (Figure S12B and Table S17B). Gene ontology and pathway enrichment analysis of these shared sets of genes showed enrichment for metabolic pathways, inflammation, haematopoiesis and neurotransmission among the DNAm and expression targets (FDR = 25%, Table S18).

To further explore the extent of shared postprandial signals, we focused on the previously identified 24 genes with differential expression postprandially (Figure 3A and Table S12). We carried out expression quantitative trait methylation (eQTM) analyses of the 24 expression signals of these genes and their DNAm levels at fasting baseline, 30m and 4h. Overall, regional DNAm-expression associations identified eQTMs for 88% of genes at nominal significance (*i.e.:* 21 of 24 genes with at least one DNAm probe associated with expression with *p* < 0.05, Figure S13). Evidence for eQTMs was observed in the peak postprandial expression signals *CPT1A, SLC25A20* and *AGPAT4.* Co-methylation patterns were also observed across multiple CpGs in genes such as the stress response mediator and mTOR inhibitor^36^ *DDIT4* and insulin and inflammation-response enhancer-binding transcription factor^37^ *CEBPD,* where neighbouring co-methylated probes showed the same direction of association with gene expression levels (Figure S14).

### Genetic regulation of postprandial DNA methylation and gene expression

Genetic variability can have substantial effects on DNAm levels. Using data from the MeQTL EPIC Database^38^ we found cis-methylation quantitative trait loci (cis-meQTLs) for 64 (49.6%) of postprandial DNAm signals, including in *PDE9A, GPT2, DAGLA, A4GALT* and *ABCC1* (Figure S15A and Table S19). Likewise, genetic variation can modulate the expression of genes and, using data from the eQTLGen phase I^39^ database, we identified cis- and trans-expression quantitative trait loci (cis- and trans-eQTLs) for 22 and 18 (92% and 75%) of our gene expression signals (Figure S15B-C and Table S20), respectively, including for *CPT1A, SLC25A20, AGPAT4, ACAA2, ADAP2, DDIT4* and *SQLE*. The circadian rhythm regulator *PER1* did not have significant eQTLs in eQTLGen phase I^39^, but had one expression quantitative trait methylation locus (eQTM) in the BIOS database (Table S21). Overall, 10 (41%) of our main gene expression signals had eQTMs (Figure S15D and Table S21), but none of the postprandial DNAm signals that were previously identified as eQTMs in BIOS.

We next sought to identify genetic variants that may affect the postprandial molecular trajectories detected. We searched for genetic regulation of DNAm at the 129 unique DMPs (in 95 unique genes) from postprandial 4h meta-analysis and trajectory signals (Table S1 and Table S6), and for genetic regulation of 24 genes with postprandial expression changes (Table S12). Using public data from the MeQTL EPIC Database^38^ and eQTLGen phase I^39^ database we identified the same variants acting as cis-meQTL and cis-eQTL for 19 genes. To explore potential links between DNAm and expression, we performed genetic colocalization analysis. We observed strong evidence for a common causal genetic variant for two CpG-gene pairs, for cg00118229–PDE9A (P(H4) > 0.98) occurring through rs2839581 (chr21:44,158,405) (Figure S16A), and for cg14880584–GPT2 (P(H4) > 0.97) occurring through rs72812428 (chr16:46,906,797) (Figure 4A). Another CpG-gene pair, cg17656260–ZNF790 (P(H4) > 0.89), was identified through colocalization, but with a credible set of 17 causal SNPs thus violating the assumption of a single causal variant and was excluded from our results. Sensitivity analysis with varying prior probabilities for cg00118229–*PDE9A* and cg14880584–*GPT2* validated our main results for these CpG-gene pairs (Figure S17). Overall, we observe evidence for genetic predisposition for coordinated DNAm and expression changes in *PDE9A,* which is implicated in lipolysis, obesity and heart disease^40–42^, and in *GPT2,* an enzyme that generates pyruvate during gluconeogenesis^26,43^. Both of these genes exhibit significant changes in DNAm levels postprandially (Figure 2; Tables S1 and S6), for example, DNAm levels in *GPT2* showed hypermethylation at 30m and 4h after test meal (Figure 4B). Therefore, the underlying genetic effects may potentially also affect the magnitude of the postprandial molecular changes observed at *PDE9A* and *GPT2*. We also observed that the genes identified from our postprandial DNAm and expression analyses (95 genes from DNAm, and 24 genes from expression analyses) harboured genetic variants that were previously identified as GWAS signals for BMI, type 2 diabetes, cholesterol levels, C-reactive protein and blood cell counts (Table S22)^44^.

**Figure 4.**
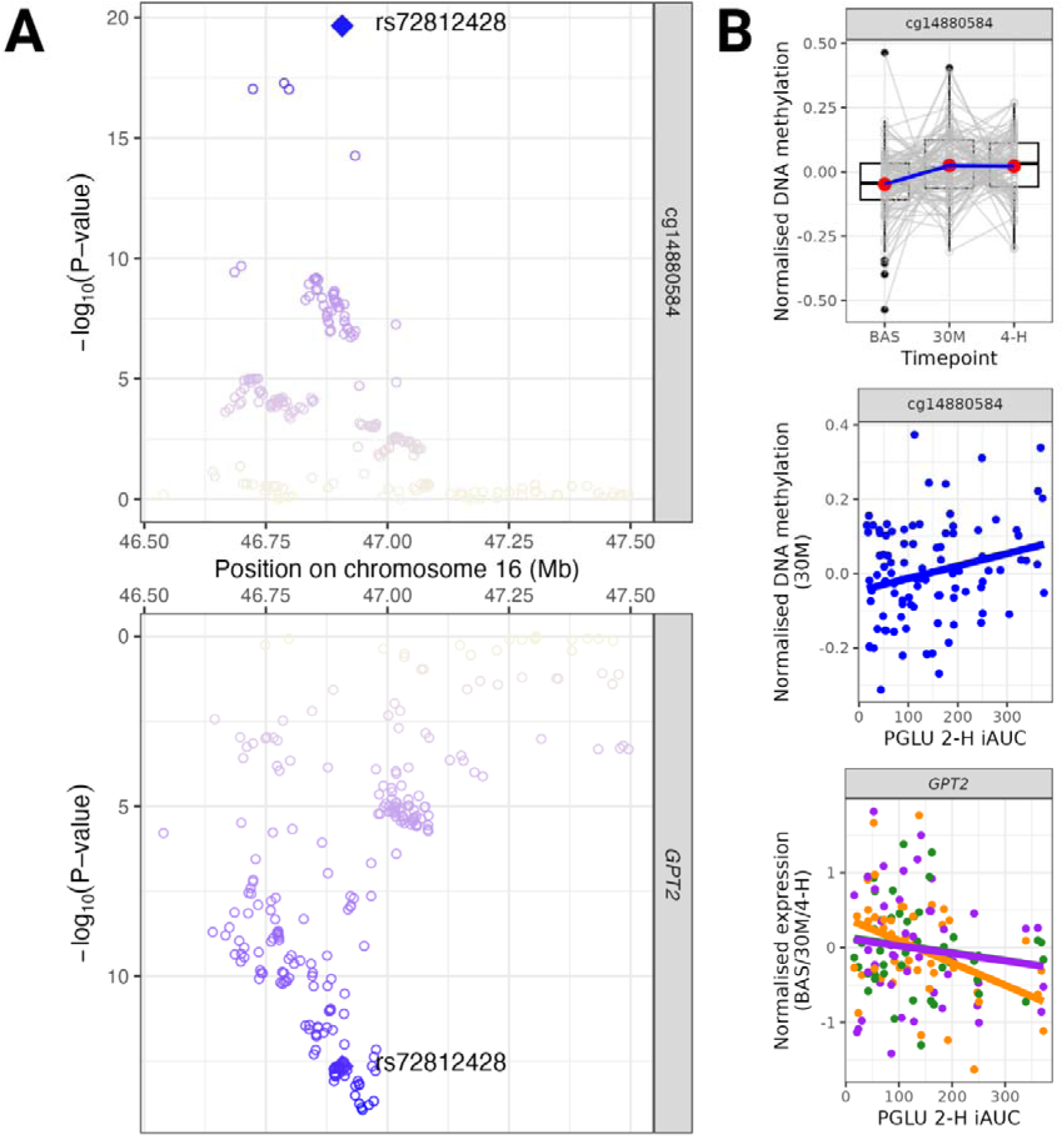
Colocalization of genetic effects on the DNAm levels of cg14880584 and expression of *GPT2*. **A.** meQTL and eQTL results from the MeQTL EPIC Database and eQTLGen phase I for the *GPT2* locus, respectively. Target SNPs for were selected within 1 Mb of cg14880584 and the zoom plot shows the location of the colocalised SNP rs72812428 (chr16:46,906,797). **B.** Postprandial trajectory of cg14880584 and associated phenotype results for cg14880584 and *GPT2*. Expression at baseline, 30m and 4h are green, orange and purple in the graph, respectively. The under-regulation of GPT2 captured at 30m after meal was overall linked to the glucose iAUC of participants (*p* < 0.05).

### DNA methylation signals of postprandial plasma glucose and triglyceride levels

The postprandial state is characterised by fast rise of plasma glucose levels within 2h after meal intake, and steady rise in blood triglycerides that peaks at 3–6h after a meal (Figure 1). Here, we link baseline and postprandial DNAm profiles to intra-individual variation in the glucose and triglycerides trajectory after a meal. We investigate if postprandial DNAm levels track inter-individual differences in the postprandial trajectories and concentrations of plasma glucose and triglycerides in females from PREDICT. We also explore if baseline DNAm levels are predictive of the postprandial metabolic response and whether baseline health can impact DNAm trajectories.

#### Postprandial DNA methylation and expression associations with plasma glucose and triglycerides

Firstly, we focused on longitudinal analyses to associate the DNAm trajectory over fasting, 30m and 4h to specific phenotypes postprandially. We observed that postprandial DNAm levels were associated with the 4h incremental area under the curve (iAUC) for triglycerides at 3 CpG sites: cg05086789 in the body of *LOC387647*, cg24013618 in the body of *GAK,* and cg21318603 in the body of *PRPSAP2* (Figure 5A and Table S23: FDR = 5%). *PRPSAP2* is a candidate gene for metabolic syndrome^45^, and its expression was also significantly associated with the overall triglycerides 4h peak rise in our data (Table S24: FDR = 5%). Specifically, cg21318603 was hypomethylated, while *PRPSAP2* was over-expressed with higher triglycerides postprandially (Figure 5B).

**Figure 5.**
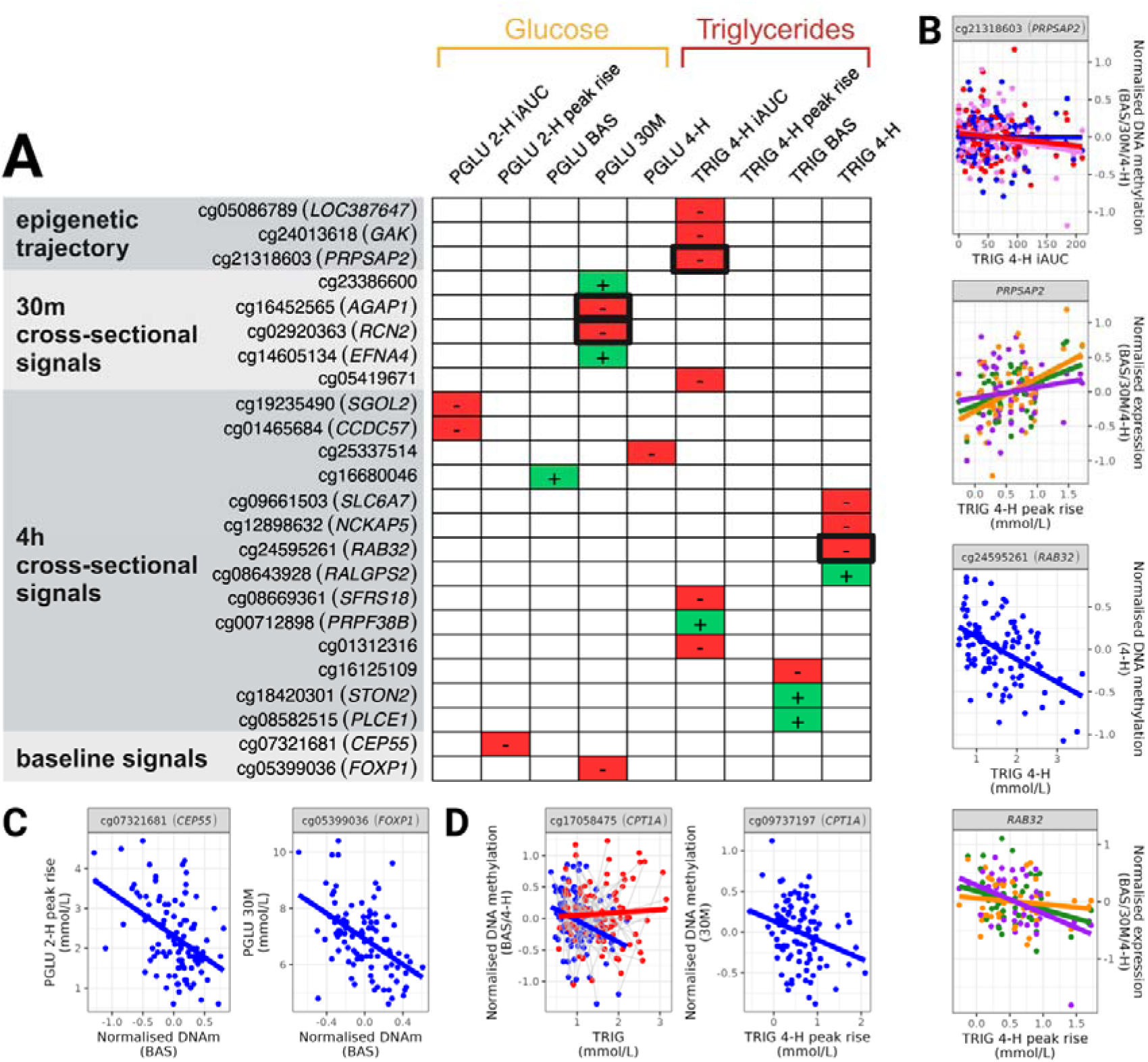
DNAm signals associated with plasma glucose and blood triglycerides after meal. **A.** Signals identified over time, cross-sectionally and at fasting baseline in association with the interindividual response to food in PREDICT (n = 104 females). Positive and negative directions of effect with the postprandial rise in glucose and triglycerides after meal are marked green and red, respectively. Signals from genes with expression changes linked to glucose and triglycerides are highlighted in bold. DNAm and gene expression signals were considered with FDR 5% and the strongest signal was cg09661503 (*SLC6A7*) for triglycerides at 4h (*p* = 3.96E-09). **B.** Examples of DNAm and gene expression links to triglycerides. Where multiple timepoints are plotted together, DNAm is blue, pink and red and gene expression is green, orange and purple for fasting baseline, 30m and 4h timepoints. **C.** Baseline DNAm signals linked to interindividual glucose levels after meal. **D.** DNAm signals from the GOLDN cohort linked to triglycerides in PREDICT (adj. *p* < 0.05). Cg17058475 (in *CPT1A*) was linked to the trajectory of triglycerides in PREDICT with effect most pronounced at baseline (baseline and 4h timepoints are blue and red, respectively).

We also carried out cross-sectional analyses linking DNAm changes to phenotype levels at specific postprandial time-points. We observed that DNAm at 30m postprandially was associated with 30m glucose, and that DNAm at 4h postprandially was most associated with postprandial lipid levels. Specifically, DNAm at 4h was associated with the concentration of triglycerides at 4h and their overall 4h peak rise at 7 DNAm signals: cg09661503 (*SLC6A7*), cg12898632 (*NCKAP5*), cg24595261 (*RAB32*), cg08643928 (*RALGPS2*), cg08669361 (*SFRS18*), cg00712898 (*PRPF38B*) and cg01312316 (Figure 5A, Table S23). Cg24595261, which was associated with triglycerides concentration at 4h, is in the extended promoter of *RAB32*. This gene is thought to play a role in lipid storage^46^, and we also detected its expression trajectory to be associated with triglycerides 4h peak rise (Figure 4B and Tables S23 and S24). Cg08643928, cg08669361 and cg00712898 are in the 5’UTR region of *RALGPS2, SFRS18* and *PRPF38B*. The expression of *RALGPS2* has been linked to cardiovascular disease^47^*, SFRS18* plays a role in fat deposition^48^ and *PRPF38B* has been found to be hypermethylated in female offspring exposed to maternal high saturated fat diet^49^. In our study *PRPF38B* was hypermethylated with the rise of blood triglycerides after meal (Table S23).

In contrast, at 30m DNAm signals captured mostly differences in glucose concentration (Figure 5A, Table S23). Four DMPs were identified for glucose concentration at 30m, including cg16452565 in the body of *AGAP1* and cg02920363 in the body of *RCN2*. Differential DNAm in *AGAP1* was previously linked to high-sugar/high-fat diet feeding^50^ and *RCN2* mediates arterial calcification in hypertension and diabetes^51^. Both of these genes were also identified in the gene expression results, linked to inter-individual differences in the glucose iAUC (Table S24). At 30m only one CpG site – cg05419671 – was associated with triglyceride levels, while at 4h postprandially 3 CpG signals were still observed to be associated with postprandial measures of glucose (Figure 5A, Table S23).

#### Baseline DNA methylation signals of postprandial glucose and triglycerides trajectories

We also tested whether baseline DNAm levels could be predictive of the postprandial metabolic response. DNAm at fasting baseline at cg05399036 (*FOXP1*) and cg07321681 (*CEP55*) was associated with the postprandial level of glucose at 30m and the overall glucose 2h peak rise, respectively (Figure 5A, Table S23). *FOXP1* controls systemic glucose homeostasis by upregulating glucose uptake in adipo- and myocytes and shutting-off glucogenesis in hepatic cells^52,53^, while *CEP55* is sensitive to glucose concentration and regulates glucose metabolism via the Akt/mTOR pathway^54^. *FOXP1* itself can be regulated by PI3K/Akt/mTOR and is newly implicated in the pathogenesis of diabetes^52,53^. Here, hypomethylation of cg05399036 and cg07321681, located in 5’UTR and TSS1500 of the *FOXP1* and *CEP55* genes, respectively, was predictive of higher glucose postprandially (Figure 5C). We did not find an association between the expression of these genes and postprandial glucose levels, but we observe an association between *FOXP1* expression levels and postprandial triglyceride levels.

Previous work by Lai et. al^5^ identified baseline DNAm levels at 5 genes – *LPP, CPT1A, APOA5, SREBF1,* and *ABCG1* – to be associated with postprandial levels of blood triglycerides in the GOLDN cohort. While we were unable to identify epigenome-wide DNAm associations for triglycerides at fasting baseline, we replicated some of Lai et. al.’s signals postprandially, specifically the hypomethylation of cg17058475 and cg09737197 in the 5’UTR region of *CPT1A* (Table S25). After multiple testing correction, cg17058475 hypomethylation was associated with the fasting levels and overall trajectory of triglycerides in blood (*b* = −0.376 ± 0.118, *p* = 8.96E-04 and *b* = −0.926 ± 0.205, *p* = 4.46E-06, respectively), and hypomethylation of cg09737197 at 30m was associated with the triglycerides 4h peak rise (*b* = −0.231 ± 0.078, *p* = 5.18E-03) (Figure 5D, Table S25). Other associations were also detected at nominal significance, including for cg00574958 (*CPT1A*) and cg11024682 in the body of the *SREBF1* gene (Table S25). In our data cg11024682 was hypermethylated, like in the GOLDN cohort, at 30m and associated with overall triglycerides 4h iAUC (*b* = 4.20E-04 ± 1.96E-04, *p* = 0.03).

Variation in human metabolic responses to food is associated with aging, where older individuals are more likely to have unfavourable glycemic^55,56^ and lipemic^57^ responses. Individuals can also be epigenetically older than their chronological age and epigenetic age acceleration (EAA) has been associated with unfavourable health outcomes and individual responses to lifestyle interventions. As such, we explored whether the participant’s EAA at fasting baseline was associated with their postprandial metabolic signature in PREDICT. EAA can be derived using different epigenetic clocks. We tested six epigenetic ageing measures, the original Horvath^58^ and Hannum^59^ clocks, which measure accelerated aging based on different sets of CpGs, PhenoAge^60^, which measures healthspan and morbidity, GrimAge^61^ and GrimAge 2^62^, which can measure both morbidity and mortality, and DunedinPACE^63^, a more recent measure for pace of aging. We tested associations between the EAA for each of these measures with postprandial levels of glucose and triglycerides, but results were not significant (Table S24). Previous research in the GOLDN cohort identified a modest association between Hannum EAA and postprandial triglycerides ^64^. We observed the same trend of effects for postprandial triglycerides and Hannum EAA in PREDICT (Table S26).

This section focused on DNAm profiles and DNAm-based molecular markers of aging at fasting baseline and evaluated their link with the postprandial metabolic response. Next, we explored whether health status at baseline relates to the postprandial molecular trajectories.

#### Baseline phenotype associations with postprandial molecular trajectories

To explore whether measures of baseline health can impact the postprandial molecular response, we analysed fasting blood concentrations of glucose and triglycerides at baseline with the postprandial DNAm levels and trajectories. We observed that fasting triglyceride concentrations showed epigenome-wide significant associations with DNAm levels at 4h postprandially at 3 signals, cg16125109, cg18420301 (*STON2*) and cg08582515 (*PLCE1*) (Figure 5A, Table S23). Fasting glucose was associated with cg16680046 DNAm levels captured at 4h postprandially as well (Figure 5A, Table S23). However, these associations were not reflected at the expression level where *STON2 and PLCE1* gene expression trajectories did not associate with baseline phenotype levels in our data (FDR = 5%).

#### Postprandial molecular changes capture variation in postprandial glucose and triglyceride levels

We assessed whether the 4h meta-analysis and postprandial trajectory results for DNAm and expression (Figures 2,3; Tables S1, S6, S12) associated with specific postprandial metabolic phenotypes. We observed that over 70% of the 129 unique DMPs from the DNAm 4h meta-analysis and trajectory analyses were also nominally associated with glucose and triglyceride levels in PREDICT (*p* < 0.05, Tables S1 and S6). This included signals cg03403093 in *INPP4A* where DNAm and triglycerides were associated at 4h (*b* = −0.059 ± 0.027, *p* = 0.022), cg26254678 in *DAGLA* where DNAm and triglycerides were linked at both timepoints including at 4h (*b* = 0.062 ± 0.022, *p* = 0.004), and cg07870237 in *ACSL6* where DNAm timepoints were overall associated with glucose peak rise (*b* = −0.041 ± 0.019, *p* = 0.043). Other nominally significant association signals included cg04924511 in *GHRL* and cg27518648 in *ABCG1* where DNAm at 30m was linked to glucose levels both at fasting and 30m after meal (*b* = −0.071 ± 0.030, *p* = 0.018 and *b* = −0.016 ± 0.008, *p* = 0.036, respectively). Nominally significant phenotype associations were also detected with gene expression changes of *INPP4A, ACSL6, GHRLOS* and *ABCG1*, as were levels of some of the main trajectory gene expression signals, such as *PER1, ACAA2, ADAP2, C15orf39, MYOF, POU2F2, SMAP2* and *AGPAT4* (*p* < 0.05, Table S12). Overall, we observed that *ABCG1* was hypermethylated at 4h (Table S1), under-expressed with the triglycerides 4h peak rise and iAUC (*b* = −0.610 ± 0.120, *p* = 5.87E-04 and *b* = −0.005 ± 0.001, *p* = 0.005, respectively), and linked to blood sugar at 30m. *ABCG1* association with postprandial glucose is in line with evidence that beyond lipid metabolism *ABCG1* participates in the metabolism of sugars by controlling the secretion and activity of insulin^19^.

Our previous results integrating genetic data into the postprandial DNAm and gene expression trajectories further highlighted two genes, *GPT2* and *PDE9A*, that were under genetic regulation (Figure 4A). Integrating these data with the phenotype results, we observe that the postprandial hypomethylation of cg00118229 (*PDE9A*) at 30m is also negatively associated to triglycerides peak rise (*b* = −0.061 ± 0.026, *p* = 0.029, Figure S16B). Similarly, we observed that postprandial hypermethylation of cg14880584 (*GPT2*) and under-expression of *GPT2* at 30m are associated with the inter-individual glucose response after meal (DNAm *b* = 3.32E-04 ± 1.33E-04, *p* = 0.026; and expression *b* = −0.002 ± 6.49E-04, *p* = 0.036) (Figure 4B).

## Discussion

In this study, we conducted the largest exploration to date of changes in blood-based DNAm and gene expression trajectories during the postprandial state in humans. We profiled blood DNAm and gene expression at fasting, and at 30m and 4h after a meal in 225 participants from two international cohorts, PREDICT and CORDIOPREV. We observed longitudinal changes in the regulation and function of key metabolic genes with implications for cardiometabolic health. We show evidence for links between DNAm and gene expression levels of certain metabolic genes, and baseline and postprandial levels of plasma glucose and triglycerides. Moreover, we identify specific scenarios of genetic regulation of coordinated changes in DNAm and gene expression, that may affect the postprandial metabolic response. The identified molecular signals could represent a response to food intake that may in turn contribute to disease risk such as postprandial rise in glucose and triglyceride levels, or alternatively these signals may reflect an adaptive response to postprandial metabolism in the organism.

The transition from a fasted to a postprandial state represents a major change in physiology that tests nutritional phenotypic flexibility of humans^65^. The metabolic processes associated with digestion and absorption of food, specifically postprandial hyperglycemia and hypertriglyceridemia, have emerged as independent risk factors for type 2 diabetes and cardiovascular diseases such as atherosclerosis and myocardial infarction^65,66^. Indeed, postprandial glucose and lipids are stronger predictors of cardiovascular events than their fasted levels^67–69^. Given the high inter-individual variability in postprandial metabolic response^1^, we aimed to characterise molecular mechanisms underlying this variability by profiling DNAm and gene expression levels at baseline and at postprandial time-points associated with the glycemic (30m) and lipemic (4h) metabolic responses. We studied participants in mid-life from two cohorts, including a sample of 104 healthy female individuals from the UK PREDICT study who were fed a high fat high carbohydrate test meal, along with a sample of 121 male diabetic coronary heart disease patients from the Spanish CORDIOPREV study who were fed a high fat meal. These differences in cohort profiles and test meal challenges highlight the robust nature of the observed epigenetic results postprandially at 4 hours, which typically corresponds to peak lipemia phase.

We observed over a hundred DNAm signals at 4h postprandially in meta-analysis of the PREDICT and CORDIOPREV studies. The majority and most-associated DNAm signals were in genes previously linked to obesity and type 2 diabetes, and lipid metabolism. These included the cholesterol efflux pump *ABCG1* and *ASIP* genes, where *ASIP* was the first obesity gene to be characterised decades ago^24^ and *ABCG1* has many roles in metabolic health, including controlling the secretion and activity of insulin and lipoprotein lipase^19^. DNAm of *ABCG1* at various CpG sites has been inversely associated with HDL cholesterol and directly associated with triglycerides in blood^5,19^. Here we report DNAm at a novel CpG site in *ABCG1* (cg27518648) where DNAm is also directly associated with postprandial lipemia at 4h. We also observed increased DNAm of *GHRL* promoter region (cg04924511) at 4h. The *GHRL* gene is responsible for synthesizing the preproghrelin precursor to ghrelin which is a hormone that acts primarily on the brain to stimulate eating and has secondary functions in the regulation of blood sugar and cellular homeostasis^22,23^. A previous postprandial DNAm study^6^ also explored *GHRL* in candidate analysis and found cg04924511 to be hypermethylated at 160m after meal intake after adjusting for cell composition, but the effect was not nominally significant. Neither we nor previous work observed a link between DNAm at cg04924511 and *GHRL* overall expression levels, but finer resolution expression analyses such as alternative splicing may need to be carried out to assess its full regulatory potential.

The additional sample time-points in PREDICT allowed us to dissect the postprandial epigenetic trajectories by also considering effects at peak glycemia (30min), as well as extending to gene expression trajectories in a sample’s subset. Postprandial trajectory analyses captured DNAm and expression changes in genes implicated in metabolism (*CAMTA1*, *CTP1A* and *SLC25A20*), inflammation (*FKBP5, PDE4D* and *CEBPD*), stress response (*CAMTA1, FKBP5*), and circadian rhythm (*PER1*). *PER1,* in addition to being a core component of the circadian clock, regulates blood pressure^70^ and is over-expressed during fasting-induced oxidative stress in the mitochondria^35^. In our case, *PER1* was under-expressed postprandially at 30m and 4h in comparison to fasting baseline, which has implications for effects of food intake as a regulator of circadian rhythm.

*CPT1A* and *SLC25A20* are both key components of lipid metabolism in mitochondrial fatty acid oxidation^71^, and ACAA2 catalyses the last step of the fatty acid beta oxidation in the mitochondria^72^. The rate of fatty acid oxidation is highest during fasting^73^, therefore *CPT1A* and *SLC25A20,* which were both under-expressed postprandially in our study, are likely downregulated after meal intake. Further lipid metabolism genes among the signals include *AGPAT4*, which catalyses a precursor to triacylglycerol^74^ and *SQLE*, which is the first oxygenation enzyme in cholesterol biosynthesis^75^. On the other hand, *PDE4D* is key in the degradation of the immunoregulatory cyclic adenosine monophosphate (cAMP) molecule^76^, while *FKBP5* binds directly to co-chaperones of the steroid receptor complex in response to stress and is under the influence of both diet-induced obesity and epigenetic regulation^77,78^.*CAMTA1* is a transcription factor that also responds to stress and is involved in insulin production and secretion^29^ with potential modest impact in type 2 diabetes^79,80^. Cg23021268 in *CAMTA1* was the most associated postprandial DNAm signal in PREDICT and was hypomethylated over the glycemic response to food at 30m*. CAMTA1* has been found to be hypermethylated after *in utero* exposure to a high-fat diet^81^, and in line with this in our results cg23021268 DNAm levels increased from 30m over the course of the lipemic response at 4h. Furthermore, multiple genes exhibited coordinated changes in postprandial DNAm and gene expression levels, including key lipid metabolism genes *ABCG1*, *CPT1A*, as well as *SLC25A20, DDIT4,* and others.

The results from postprandial 4h meta-analysis and DNAm and expression trajectories in PREDICT females alone identified 119 unique genes where molecular genomic changes were identified postprandially. The genes were enriched for pathways involved in fat metabolism, inflammation, haematopoiesis and neurotransmission. We successfully replicated a proportion of these DNAm and gene expression postprandial trajectory results in two independent studies ^6,14^. The replication studies were carried out in samples from different populations and at different age ranges, with different test meal challenges, and in males only or mixed male and female samples who were sampled at different time-points postprandially, and in some cases profiled using different molecular profiling technologies. These differences highlight the robust nature of the replicated postprandial blood changes. The first replication study by Rask-Andersen et al.^6^ analysed whole blood DNAm using Illumina Infinium HumanMethylation450 BeadChip and microarray gene expression in 26 healthy men before and 160m after a standardized mixed breakfast of whole wheat, fruit and cheese. The second replication study by GOTO^14^ collected transcriptomic data from 85 healthy older participants before and 30m after a standardized meal replacement shake. In our study we replicated 15% of DMPs reported by Rask-Andersen et al.^6^ and identified 7 genes with postprandial gene expression changes across PREDICT, Rask-Andersen et al.^6^ and the GOTO^14^ studies: *CPT1A, SLC25A20*, *PER1, ACAA2, DDIT4, C15orf39* and *MYOF.* These results demonstrate the robust nature of postprandial gene expression changes at these seven genes involved in lipid metabolism, circadian rhythm and cellular responses to stress and inflammation.

We next observed associations between some of the postprandial DNAm and expression signals with postprandial levels of glucose and triglycerides. Cross-sectionally, DNAm at 30m and 4h was most strongly associated with glucose and triglyceride levels, respectively. This was expected since glucose peaks first, and chylomicron triglycerides and VLDL triglycerides only peak at 3–4h or 4–6h after meal^82^, respectively. The genes that harbour the signals that underlie these associations included *RAB32* (linked to lipid storage^46^, also detected with gene expression), *RALGPS2* (linked to cardiovascular disease^47^)*, SFRS18* (fat deposition^48^), and *PRPF38B* (linked to high saturated fat diet^49^). Longitudinally, the postprandial DNAm trajectories of several genes including *PRPSAP2* (metabolic syndrome^45^) associated with the postprandial triglyceride levels, measured by the iAUC. In contrast, baseline levels of DNAm were associated with postprandial levels of glucose, rather than triglycerides. Fasting DNAm levels in two genes, *FOXP1* (controls systemic glucose homeostasis ^52,53^) and *CEP55* (regulates glucose metabolism via the Akt/mTOR pathway^54^) were linked to postprandial glucose with implication for glucose homeostasis and metabolism. This result is intriguing as it has implications for the potential of fasting DNAm levels as a baseline prediction tool of the postprandial glucose peak, which is a risk factor for cardiometabolic health. Lack of similar datasets limited our ability to pursue replication of our postprandial glucose and triglyceride DNAm associations, but we explored how previous findings replicated in our data. Specifically, the GOLDN study profiled fasting baseline DNAm levels in CD4+ T-cells in 979 subjects of European descent, and identified associations with postprandial lipid levels after a high fat meal^5^ in 5 genes, *CPT1A, ABCG1, SREBF1, LPP* and *APOA5*. We replicated triglyceride associations with DNAm at cg17058475 and cg09737197 in *CPT1A* after multiple testing correction, and at cg11024682 in *SREBF1* at nominal significance, albeit not at baseline. Particularly, in our data postprandial DNAm in cg09737197 was inversely associated with the triglycerides peak rise, which controls for baseline triglycerides by subtracting the fasting levels from their 4h peak. Furthermore, signals in *ABCG1* and *LPP* were hypermethylated with triglycerides postprandially in the GOLDN dataset, and while we did not replicate these, we observed that *ABCG1* and *LPP* were under-expressed with triglycerides, particularly with stronger triglycerides peak rises (*p* < 0.001) (Table S27).

The 119 unique genes identified from postprandial changes in DNAm and expression were enriched for pathways related to fat metabolism, inflammation, haematopoiesis and neurotransmission. Integration of these results with the EWAS Catalog, identified enrichment of previously published associations of these CpGs with traits such as BMI, C-reactive protein and blood cell lineage-specific proteins. Using publicly available databases we assessed whether genetic background may impact the identified postprandial DNAm and expression signals. We identified cis-meQTLs and cis-eQTLs for 49.6% and 92% of our epigenetic and expression signals (Tables S19 and S20), respectively. In the GWAS Catalog, multiple genetic variants within the genes identified in our study were previously associated with traits such as type 2 diabetes, cholesterol, and also BMI, C-reactive protein and blood cell counts. Overall, there is strong prior evidence that human genetics plays a role in determining the levels of DNAm and expression of genes. Furthermore, our colocalization analyses found evidence for shared genetic regulation of DNAm and gene expression levels at two genes with postprandial DNAm and expression changes, *PDE9A* (linked to lipolysis, obesity and heart disease^40–42^) and gluconeogenesis^26,43^ gene *GPT2*. The signal in gluconeogenesis gene *GPT2* was hypermethylated postprandially, and at 30m its hypermethylation and under-expression was also associated with postprandial glucose response. Taken together these results provide specific examples of potential genetic regulation of the postprandial molecular genetic response that may be mediated *via* DNAm and gene expression effects, and in the case of *GPT2* strongly relate to postprandial glucose levels.

We considered biological scenarios that may explain the observed changes in postprandial DNAm levels as early as 30m after meal intake. First, we considered mammalian DNA methyltransferase kinetics. *In vitro* experiments by Adam et. al ^83^ showed that murine DNMT1 can methylate up to 70% of DNA within the first 30m of incubation with hemimethylated DNA substrate. The genomic context of CpG sites matters for timing of DNAm and Adam et. al found that demethylation is favoured at sites where DNMT1 has less affinity for the DNA molecule^83^. Timing of demethylation has been further explored by Ravichandran et al.^84^ where authors showed that mammalian TET3 can demethylate up to half of CpG sites in the canonical E-box ‘CACGTG’ motif within 30 minutes of *in vitro* incubation. Given the kinetics of mammalian DNA methyltransferase and de-methylation processes, although it is possible that the detected effects represent novel methylation and demethylation events, another plausible scenario is postprandial change in blood cell composition. In line with this, we detected changes in blood cell subtypes following meal intake, most prominent in the proportions of monocytes, granulocytes and natural killer cells (Figure S4). Changes in blood cell proportion may be attributed to overproduction of particular cell sub-populations, or sequestration, or changes in activation of specific cell types. Indeed, previous studies have reported changes in postprandial blood cell proportions ^6^ and evidence for postprandial activation of leukocytes^85^. For example, in healthy subjects, postprandial studies following fat intake have reported an increase in neutrophil counts around peak postprandial lipemia, as well as increased activation of monocytes and neutrophils^86^. A specific example of postprandial leukocyte activation that is likely relevant to our findings is postprandial activation of monocytes, such as lipid-laden monocytes. Khan et al.^86^ reported that in comparison to controls, obese subjects with metabolic syndrome had both higher fasting levels of lipid-laden foamy monocytes and significant postprandial increase in activation markers on these nonclassical monocytes. Activated monocytes with greater inflammatory and adhesion potential may lead to increased transendothelial migration, contributing to progression of atherosclerosis.

One of the main limitations of our study is that analyses were conducted in blood, a heterogeneous collection of cells. DNAm in PREDICT was generated for whole blood samples, while in CORDIOPREV the polymorphonuclear leukocyte blood fraction was extracted. Therefore, the meta-analysis signals that we identify in postprandial lipemia are relevant only to polymorphonuclear leukocytes and changes in these cells. Previous work by Rask-Andersen et al. ^6^ highlighted the importance of accounting for changes in blood cell composition when considering postprandial DNAm. We correct for blood cell type heterogeneity by including estimates of blood cell proportions as covariates in the linear associations. However, the level of resolution of the estimated blood cell types is not high, for example, our data do not differentiate between DNAm signals in monocytes and in lipid-laden foamy monocytes. We attempted to explore this heterogeneity further by profiling DNAm in a small sample of matched CD4+ T-cells and whole blood, where we see overall some consistency in DNAm levels, with 41% of tested signals showed the same direction of effects. However, our study design did not target additional blood cell subtypes, such as monocytes, granulocytes and natural killer cells, which should be explored in future studies of postprandial molecular genomic trajectories.

Circadian rhythm and seasons can affect human metabolism^87,88^ and thus may impact postprandial responses to food. To this end, we accounted for seasonality as a covariate in the linear association models. All participants in PREDICT and CORDIOPREV had the test meal in the morning after an overnight fast, therefore minimising potential effects of circadian rhythm variability on DNAm levels. To further explore potential effects of circadian rhythm, we interrogated a previously published dataset of CpGs with reported circadian oscillation in human DNAm^33^ but we did not find evidence for a major effect of time of day. Another limitation is that many analyses in PREDICT included only females and it is highly likely that females and males yield different DNAm and expression responses to a food challenge, as previously observed for gene expression^14^. Because the CORDIOPREV sample was male, our PREDICT-CORDIOPREV meta-analysis identified sex-shared effects. Sex-specific analyses were challenging to undertake in our study due to confounding with differences in phenotypes (the PREDICT sample was composed of healthy middle-aged British females while the CORDIOPREV study was composed of middle-aged Spanish coronary heart disease patients) and test meal challenges (PREDICT meal was high in carbohydrates and fat while the CORDIOPREV meal was high in fat alone, with different food composition). Lastly, the CORDIOPREV samples were available only fasting baseline and 4h, therefore, giving us greater power to identify DNAm signals associated postprandially at 4h during peak lipemia, than during peak glycemia. Replication of the gene expression results from PREDICT in the GOTO study also suffered from similar limitations due to differences in study population, study design, meal composition and timepoints availability. Due to these reasons, interpreting the directions of effect of signals postprandially across studies was challenging. Future postprandial analyses of DNAm and gene expression will also benefit from time-matched larger replication samples to ascertain impact of food response in cardiovascular disease.

## Conclusion

Postprandial changes in blood DNAm and gene expression were observed at key metabolic genes postprandially, from baseline fasting to 30m and 4h after meal intake. Some of the signals reflect the inter-individual variation observed in blood glucose and triglyceride levels after a meal, with potential to act as either mediators of, or signals reflecting postprandial metabolism processes. Overall our findings identify postprandial molecular genetic changes in genes related to metabolism, inflammation, circadian rhythm and other processes, and emphasize the need for future studies into postprandial molecular changes within specific cell sub-types. The results highlight DNAm as a key regulatory mechanism of gene expression over the human postprandial state, with implications in cardiometabolic health outcomes.

## Materials and Methods

### Study samples

In this study we used blood samples from participants recruited in the ZOE PREDICT 1 (PREDICT)^15^ and CORDIOPREV^16^ studies. Both studies performed a food challenge on day one of clinical visit and blood samples were extracted after overnight fast, and postprandially after food intake. The PREDICT food challenge aimed to stress the body’s overall metabolic response to food in 1002 healthy middle-aged participants from the UK (mean age = 45.6 ± 11.9 years, BMI = 25.6 ± 5.0 kg/m^2^). Participants fasted overnight and following baseline blood draw participants were given breakfast (muffin and milkshake) that was high in carbohydrates and fat (850 kcal from 85g of carbohydrates, 50g of fat and 15g of protein). In addition to fasting, blood was drawn at 15, 30, 60, 120, 180, 240m after test meal for analysis of the postprandial response. The CORDIOPREV food challenge aimed to study the body’s response to dietary fat in 1002 middle-aged coronary disease patients from Spain (mean age = 59.5 ± 0.2 years, BMI = 31.1 ± 0.1 kg/m^2^). A fasting blood sample was taken immediately afterwards the patients ingested a weight-adjusted meal (0.7 g fat and 5 mg cholesterol per kg body weight) with 12% saturated fatty acids (SFAs), 10% polyunsaturated fatty acids, 43% monounsaturated fatty acids, 10% protein and 25% carbohydrates from olive oil, skimmed milk, white bread, cooked egg yolks, and tomatoes. In addition to the fasting sample, blood was extracted every hour during the 4 h after test meal, according to established protocols. Both PREDICT and CORDIOPREV test meals were prepared by nutritionists and volunteers rested and consumed no food for 4h but were allowed to drink water. Here, we used samples collected at fasting and 30m and 4h postprandially in the PREDICT study to explore the DNAm and expression trajectories associated with the glycemic and lipemic responses to food. Additionally, we used blood samples collected at fasting and 4h postprandially in CORDIOPREV to meta-analyse the longitudinal DNAm changes associated with postprandial lipemia in PREDICT and CORDIOPREV.

### Extraction and processing of biological data

#### DNA methylomes

DNAm profiles were extracted from 360 whole blood samples collected at 3 timepoints from 120 participants in PREDICT and 292 polymorphonuclear leukocyte blood fraction samples collected at 2 timepoints from 150 diabetic coronary heart disease patients in CORDIOPREV. An additional 30 DNAm profiles were extracted from CD4+ T cell samples collected at 3 timepoints from 10 participants in PREDICT for blood cell type analysis. Genomic DNA was extracted and bisulphite converted using the EZ-96 DNA Methylation Kit (Zymo Research, Orange, CA, USA). Subsequent methylation analysis was performed on an Illumina (San Diego, CA, USA) iScan platform using the Infinium MethylationEPIC BeadChipv1 according to standard protocols provided by Illumina. GenomeStudio software version 2011.1 with Methylation Module version 1.9.0 was used for initial quality control of assay performance and for generation of methylation data export files.

DNAm data was processed using R Bioconductor software^89^ and methods previously described^4^. First, samples with median methylated and unmethylated signal ratio < 10.5 were excluded using the *minfi* package^90^. Second, background correction, dye bias correction and quantile normalization were performed with the ENmix package^91^. Underperforming probes and outlier samples were identified with standard parameters and signals with detP > 0.000001 and nbead < 3 were excluded. DNAm beta values were estimated after applying the Regression on Correlated Probes method^92^ in ENmix, and subsequently converted to DNAm M-values using the *lumi* package^93^. Maximum missingness threshold in samples and probes was 1% and 5%, respectively, and polymorphic or probes that mapped to multiple locations in the genome were removed. A total of 356 and 279 whole blood samples from 120 and 148 participants, and 755,438 and 747,764 autosomal probes, were kept in PREDICT and CORDIOPREV data, respectively. Blood cell proportions of samples were estimated using the Houseman method^94^ as implemented in Horvath’s DNA Methylation Age Calculator^58^.

#### Transcriptomes

Bulk RNA was extracted from 150 whole blood samples collected at 3 timepoints from a subset of 50 female participants in PREDICT. Extractions and RNA sequencing were performed by Novogene (Cambridge, UK) using Illumina’s Novaseq 6000 instrument.

Quality control of sequencing reads was performed using FastQC^95^. FastQScreen^96^ was used to assess contamination of samples. Afterwards, paired reads were adaptor and quality trimmed with Trim Galore (https://www.bioinformatics.babraham.ac.uk/projects/trim_galore/), which implements trimming with cutadapt^97^, with minimum 3 bp adaptor overlap and Phred score of 20. The STAR aligner^98^ was used to map sequencing reads to the human genome GRCh38 build, with the Y chromosome masked, 2-pass mapping and standard parameters. Alignments were then filtered with SAMtools^99^ and only uniquely mapped properly paired reads were kept for downstream analyses. Featurecounts^100^ was used to annotate and quantify genes using the ENSEMBL annotation of the reference genome (version GRCh38.92) and standard parameters. Sample normalisation was performed using the transcripts per million (TPM) method and genes were kept if they were expressed at ≥0.5 TPM in at least 10% of samples. For each gene, TPMs were further rank-based inverse normalised prior to association analyses. All 150 samples from the 50 participants and 23,466 autosomal gene-level transcripts were kept after data processing.

### Statistical analyses

The PREDICT and CORDIOPREV DNAm samples were 87% female and 84% male, respectively, and we analysed PREDICT females and CODIOPREV males alone in this study. Altogether, and after excluding samples missing the covariates that we integrated into the models, we analysed whole blood DNAm in 308 samples from 104 PREDICT females, 232 samples from 121 CORDIOPREV males, and expression in a time-matched subset of 148 samples from 50 females in PREDICT.

#### Meta-analysis of PREDICT and CORDIOPREV DNAm signals

We used the PREDICT and CORDIOPREV samples to meta-analyse the longitudinal DNAm changes associated with postprandial lipemia at 4h. For this we applied linear mixed-effects models using the lme4 package^101^ with DNAm M-values as the response to timepoint coded as factor 0/1 for fasting-4h time-point differences. We adjusted models for age, BMI, smoking (using 0 for never smoker, 1 for ex-smoker and 2 for current smoker), estimated blood cell proportions (CD8T, CD4T, NK, B cells, granulocytes and monocytes) and season (coded as 0/1/2/3 for Spring/Summer/Autumn/Winter) as fixed effects, chip, chip position and the participant as random effects, and family and zygosity in PREDICT as random effects as well. METAL^102^ was used for fixed-effects inverse variance weighted meta-analysis of 746,953 autosomal probes in PREDICT and CORDIOPREV. Results are presented after multiple testing with false discovery rate (FDR) = 5% and filtered for heterogeneity with HetISq < 85% and HetPval ≥ 0.01. Inflation in CORDIOPREV and PREDICT results was 1.3 and 1.1, respectively, which we accepted as maximum reasonable inflation in our analyses considering our limited sample sizes and the overall metabolic changes that can occur after meal. Sensitivity analyses were performed using raw DNAm β-values as opposed to M-values, as well as using the complete sample of mixed female and male participants with all covariates in each cohort (104 females and 16 males in PREDICT; 121 males and 24 females in CORDIOPREV).

#### Analysis of DNA methylation and expression trajectories in PREDICT

For analysis of DNAm and expression trajectories we used the PREDICT sample that contained timepoints at fasting and postprandially at 30m and 4h. For DNAm analysis: DNAm M-values were the response to timepoint coded as factor 0/1/2 for fasting-30m-4h differences in a linear mixed effects model adjusted as immediately above. For expression analysis: gene-level inverse normalised TPMs were the response to timepoint coded as factor 0/1/2 in a linear regression model. The model for expression was adjusted for PC2-12, GC mean and RIN. PC1 was not included in the model since it was correlated with the exposure variable, timepoint. PC2-12 explained 31% of total variance in the data and were correlated with covariates such as age, BMI, smoking, blood cell counts, and technical covariates such as rack. GC mean and RIN were added separately to the model because they were highly correlated with PC1. Separately, we trialled gene expression analysis with a linear mixed effects model adjusted similarly to our DNAm analyses and including the transcriptomic-specific covariates. Results from this model were very inflated (λ = 4.1 vs λ = 1.1 in model with PCs). As such, we present results from the linear model adjusted with PCs first and use the linear mixed effects model results for sensitivity analysis. Post-hoc analysis of timepoints was performed using DNAm M-value and gene expression residuals with pairwise t-tests across the 3 possible groups (0-30m, 0-4h and 30m-4h DNAm). The Holm method was used to correct *p*-values across the groups (adj. *p* < 0.05).

#### Expression quantitative trait methylation analysis

eQTM analyses were carried for the 24 gene expression signals identified from the main gene expression analysis. Coordinated changes at fasting baseline, 30m and 4h were explored across 146 time-matched samples using DNAm M-values as the response variable to residuals of gene expression so that we could adjust DNAm directly in the linear model. Covariates for DNAm were the same as previously, and gene expression residuals reflected normalised expression adjusted for PC2-12, RIN and GC mean. Associations are reported with nominal significance (*p* < 0.05) within target gene. Finally, we inspected gene co-methylation levels and correlation between methylated probes and gene expression and report correlation scores computed using Pearson’s correlation coefficient.

#### Analysis of DNAm and gene expression links to glucose and triglycerides

Genome-wide DNAm and expression associations were performed with postprandial glucose and triglyceride levels in PREDICT (Table S28). To track changes over the postprandial phase we used the 2h peak rise and 2h iAUC, and 4h peak rise and 4h iAUC, for plasma glucose and triglycerides, respectively. The peak rise metric, particularly, adjusts for baseline levels by subtracting the fasting glucose and triglyceride levels from their postprandial peak. In addition to this, we tested associations with direct plasma glucose at fasting, 30m and 4h, and triglycerides at fasting and 4h. Blood triglycerides were not profiled at 30m in PREDICT and 30m triglycerides are not associated with postprandial lipemia. Measurement outliers were excluded from the analysis by removing concentration values that were larger or smaller than 3 standard deviations from the mean.

First, we tracked the trajectory of DNAm and expression with the trajectory of plasma glucose and triglycerides. For this, we used DNAm and expression at fasting, 30m and 4h like in previous models plus timepoint and the interaction of timepoint and glucose or triglycerides as additional covariates. In expression analyses PC3, and PC7 and PC8, were excluded due to correlation with triglycerides and glucose, respectively. Beyond this analysis, we also used the 3-timepoint epigenetic and transcriptomic data in overall associations with the glucose and trigylceride peak rises and iAUCs.

Second, we performed cross-sectional analyses of DNAm at either 30m or 4h. DNAm residuals adjusted for timepoint and covariates were used due to the smaller number of samples included cross-sectionally (≈ one third of the original sample size). We tested DNAm associations with glucose at fasting, 30m and 4h, triglycerides at fasting and 4h, and their overall peak rises and iAUCs. Linear regression models were employed in this case. Thirty-minute and 4-hour DNAm associations with fasted glucose and triglycerides were performed to investigate role of participant health in determining postprandial DNAm levels.

Finally, we tested associations between fasting DNAm and postprandial glucose and triglycerides. For this, we used linear models where residuals of DNAm at fasting baseline were the predictors of changes in glucose and triglycerides at fasting, 30m, and 4h, as well as their overall peak rises and iAUCs. In addition, we used DNAm at 30m to predict 4h glucose and 4h triglycerides.

DNAm and expression associations with the phenotypes were overall inflated and Bacon correction^103^ of *p*-values was employed. Despite this, gene expression results remained overwhelmingly inflated (highest λ = 2.3 after Bacon) so we used them only to cross-validate findings from DNAm in this study. Results are reported after multiple testing FDR = 5%, and signals from the main postprandial analyses, *i.e.:* timepoint analyses (Tables S1, S6 and S12), that have nominal significance with the phenotypes are also reported (*p* < 0.05). Cross-sectional and baseline gene expression are not reported due limited number of samples available to perform the analyses (expression n_max_ = 50 samples per timepoint).

#### Analysis of epigenetic aging link to postprandial glucose and triglycerides

The epigenetic age acceleration (EAA) of participants at fasting baseline was used to predict changes in the blood glucose and triglycerides of participants after meal. For this we used 6 variables of EAA – AgeAccelerationResidual, AgeAccelerationResidualHannum, AgeAccelPheno, AgeAccelGrim, AgeAccelGrim 2 and DunedinPACE – in association with glucose and triglyceride levels at fasting, glucose levels at 30m postprandially, triglyceride levels at 4h postprandially, glucose 2h peak rise and iAUC, and triglycerides 4h peak rise and iAUC. Linear mixed models were adjusted for the same covariates used in the main postprandial analyses except for age at visit since EAA already accounts for chronological age. All EAAs were calculated using Horvath’s calculator^58^ apart from DunedinPACE ^63^, where we used the DunedinPACE R package (https://github.com/danbelsky/DunedinPACE) with standard parameter values, and GrimAge 2^62^.

#### Replication of DNAm and gene expression signals

The GOLDN study^5^ associated DNAm at baseline with postprandial triglycerides, while Rask-Andersen et al.^6^ profiled DNAm and expression postprandially at 160m to study changes in relation to fasting. Here, we used previously published data from Rask-Andersen et al. and the GOLDN cohort to perform a replication analysis of their findings in our trajectory and triglycerides associations, respectively. The GOTO study^14^ collected transcriptomes at fasting baseline and 30m postprandially, pre- and post-combined nutritional and physical intervention, and we replicated our expression trajectory results in this cohort. Replication results are presented after Bonferroni correction of *p*-values (*p* = 0.05/no. of probes or genes).

### Exploration of changes to postprandial blood cell composition

After findings by Rask-Andersen et al.^6^ we evaluated whether the postprandial DNAm signals captured in our analyses were being driven by changes in blood composition after meal. For this, we performed Wilcoxon signed ranks tests (and the Kruskal-Wallis test in PREDICT) to capture overall changes in blood using the Houseman et al. ^94^ blood cell count proportions estimated in our study. CD4+ T cell DNAm profiles were also available for a subset of mixed-sex participants in PREDICT. As such, we compared the directions of effect from our main postprandial lipemia analysis to the effects observed at 4h in CD4+ T cells. For this, we used raw DNAm β-values from a matched 10-participant subsample with whole blood and CD4+ T cells available. We tested changes in 27 CpGs showing the same directions of effect before and after adjustment for covariates in PREDICT, and report the directions of effect across observed in whole blood and CD4+ T cells, as well as their overall DNAm trajectory in our study.

### Gene ontology and pathway enrichment analysis

The peak 200 genes with the lowest P-values from our main DNAm and expression results were selected for term enrichment analysis with Enrichr^104^. Enrichment was performed across 5 databases, i.e.: GO Biological Process (v. 2023), GO Molecular Function (v. 2023), KEGG (v. 2021), Reactome (v. 2022) and Wikipathway (v. 2023). Background gene lists comprised all autosomal genes analysed in this study: 25,269 genes, 25,292 genes and 23,042 genes from the 4h DNAm meta-analysis, DNAm trajectory analysis and expression trajectory analysis, respectively. Only signals with low heterogeneity were included among the peak 200 signals from the 4h DNAm meta-analysis results. Significance of term enrichment was assessed with FDR = 5%.

Common genes from both DNAm and expression results were selected at a nominal P-value (*p* < 0.05) and used for term enrichment here as well. Enrichment analysis was performed with Enrichr and the significance of terms was considered with FDR = 25%. In this case the background was comprised of only genes present in both the expression and DNAm datasets.

### EWAS and GWAS Catalog look-up

The peak 200 signals from the 4h epigenetic meta-analysis and epigenetic trajectory results were searched in the EWAS Catalog^28^ to perform term enrichment analysis. For this, traits with similar or redundant names were merged, Fisher’s exact tests were conducted, and significant enrichment of terms is reported with FDR = 5%. Second, genes that passed the multiple testing threshold (FDR = 5%) in our main DNAm and gene expression analysis were searched in the GWAS Catalog^44^ and the terms of genes with SNPs are reported.

### Cross-reference of signals against genetic effects

CpGs from the main epigenetic results of this study (Tables S1 and S6) were cross-referenced against the MeQTL EPIC Database (https://epicmeqtl.kcl.ac.uk/)^38^ to determine the genetic effects on the DNAm signals reported. Similarly, genes from the main expression trajectory results (Table S12) were cross-referenced against the eQTLGen phase I database^39^ to evaluate genetic effects on the expression signals reported. DNAm effects on expression (eQTM) were searched in the BIOS database^105^. MeQTLs, eQTLs and eQTMs were considered at the threshold reported by the authors (FDR = 5% across databases). Only cis-meQTLs and cis-eQTMs from MeQTL EPIC Database and BIOS were considered while both cis and trans-eQTLs from eQTLGen phase I are reported.

### Genetic colocalization analysis

A total of 18 CpG–gene pairs with the same meQTL/eQTL SNP were identified from the MeQTL EPIC Database (https://epicmeqtl.kcl.ac.uk/)^38^ and eQTLGen phase I database^39^. Afterwards genetic colocalization analysis was performed to search evidence of genetic regulation of postprandial DNAm and gene expression. Bayesian genetic colocalization was performed according to Giambartolomei et al.^106^. For this, cis-meQTLs from the MeQTL EPIC Database were selected within 1 Mbp of target CpGs and matched cis-eQTL SNPs were selected from the eQTLGen Phase I summary statistics. Genetic variants from both datasets were harmonised, and SNPs with more than 0.1 difference in estimated minor allele frequency between the two studies were discarded. Prior probabilities for the SNPs were estimated and colocalization was performed assuming a single causal variant, using the R package coloc^106^ and the function *coloc.abf*. CpG–gene pairs were considered to have strong evidence of colocalization if the posterior probability was higher than 0.8 (*i.e.:* P(H4) > 0.80). Sensitivity analysis was performed to validate the robustness of the results. For this, we used the *coloc.abf* function with the default prior probabilities (*p*_1_ = *p*_2_ = 1E-04*, p*_12_ = 1E-05), and then used the sensitivity function to search evidence for colocalization while varying *p*_12_ values. Further details on the CpG-gene pairs tested and the estimation of priors is included in the Supplementary Information file.

## Acknowledgements

We thank the participants and clinical teams of the PREDICT, TwinsUK, CORDIOPREV and GOTO studies. We thank the the Córdoba branch of the Biobank of the Sistema Sanitario Público de Andalucía (Andalusia, Spain) for providing the human biological samples for CORDIOPREV. We would also like to thank the EASP (Escuela Andaluza de Salud Publica), Granada, Spain, which performed the randomization process for CORDIOPREV.

We acknowledge use of King’s Computational Research, Engineering and Technology Environment (CREATE, https://doi.org/10.18742/rnvf-m076) in this work.

## Author contributions

JTB, SEB and JMO designed the study. JTB oversaw the study with inputs from JMO and SEB. RC led the analysis. LDR, JMO, JFA-D, OR-Z and JL-M contributed to data acquisition. MW oversaw epigenetic profiling. TG, FB, SV, LS, YR, MT, CC, LDR, MW, PES contributed to data processing and analysis, and results interpretation. KSS, BTH and TDS contributed to results interpretation. RC and JTB wrote the manuscript. RC and SV prepared visualisation of results. All authors reviewed and approved the manuscript.

## Declarations

### Ethics approval

**PREDICT/TwinsUK:** Ethical approval was granted by the National Research Ethics Service London-Westminster, the St Thomas’ Hospital Research Ethics Committee (EC04/015 and 07/H0802/84). All research participants have signed informed consent prior to taking part in any research activities.

**CORDIOPREV:** Ethical approval was granted by the Ethics Committee from Reina Sofia University Hospital (No. 1496/27/03/2009), following the principles of the Helsinki Declaration and good clinical practices. All research participants have signed informed consent prior to taking part in any research activities. The study is registered in Clinicaltrials.gov (NTC00924937).

### Consent for publication

All authors have read and approved the manuscript for publication.

## Data availability

The epigenetic and transcriptomic data from PREDICT/TwinsUK is available through EGA [ID] for DNAm and EGA [ID] for gene expression. Additional PREDICT/TwinsUK individual-level data are not permitted to be shared or deposited due to the original consent given at the time of data collection. However, access to these data can be applied for through the TwinsUK data access committee. For information on access and how to apply http://twinsuk.ac.uk/resources-for-researchers/access-our-data/.

In relation to the epigenetic data from CORDIOPREV Study, collaborations with the study are open to Biomedical Institutions, always after an accepted proposal for scientific work. Depending on the nature of the collaboration, electronic data, hard copy data, or biological samples should be provided. All collaborations will be made after a collaboration agreement. Terms of the collaboration agreement will be specific for each collaboration, and the extent of the shared documentation (i.e., deidentified participant data, data dictionary, biological samples, hard copy, or other specified data sets) will be also specifically set on the light of each work.

## Supplementary Information

**Figure S1.** Alternative plot of DNAm signals identified at 4h postprandially in PREDICT and CORDIOPREV (n = 225 participants). The effect sizes displayed here are based on DNAm β-values instead of M-values (**Figure 2A** from the main text).

**Figure S2.** Postprandial DNAm trajectories of the 108 DMPs identified in the 4h epigenetic meta-analysis. Changes were analysed at fasting baseline (BAS), 30m (30M) and 4h (4-H) in PREDICT. Normalised DNAm levels are DNAm M-values residualised for potential confounders.

**Figure S3.** Postprandial DNAm trajectories of the 108 DMPs identified in the 4h epigenetic meta-analysis. Changes were analysed at fasting baseline (BAS), 30m (30M) and 4h (4-H) in PREDICT. DNAm levels are raw unadjusted DNAm β-values.

**Figure S4.** Postprandial blood cell proportion changes in the PREDICT (**A**) and COPRDIOPEV (**B**) studies. Changes were analysed at fasting baseline (BAS), 30m (30M) and 4h (4-H) after meal using the Wilcoxon signed rank test for each pair of samples and the Kruskal-Wallis test for overall differences in PREDICT. Estimated blood cell proportions are based on DNAm patterns from whole blood in PREDICT and polymorphonuclear leukocyte blood fractions in CORDIOPREV.

**Figure S5.** Directions of effect at 4h postprandially using M- or β-values as the response variable to timepoint in the PREDICT and CORDIOPREV cohorts (n = 225 participants). Signals from the main postprandial DNAm 4h meta-analysis (**Figure 2A; Table S1**) are highlighted in blue.

**Figure S6.** Postprandial DNAm trajectories of the 27 DMPs identified in the main epigenetic trajectory analysis. Changes were analysed at fasting baseline (BAS), 30m (30M) and 4h (4-H) in PREDICT. Normalised DNAm levels are DNAm M-values residualised for potential confounders.

**Figure S7.** Postprandial DNAm trajectories of the 27 DMPs identified in the main epigenetic trajectory analysis. Changes were analysed at fasting baseline (BAS), 30m (30M) and 4h (4-H) in PREDICT. DNAm levels are raw unadjusted DNAm β-values.

**Figure S8.** Postprandial DNAm trajectories in CD4+ T cells (CD4T) and matched whole blood (WB) samples. Trajectories are represented for signals from the main postprandial lipemia meta-analysis (**Figure 1, Table S1**) with consistent directions of effect in the whole blood raw DNAm β-values from the 10-participant subset with CD4+ T cells available.

**Figure S9.** Volcano plots of gene expression signals identified at 30m (left) and 4h (right) postprandially in PREDICT (n = 50 participants).

**Figure S10.** Postprandial expression trajectories of the 24 genes identified in the main expression trajectory analysis. Changes were analysed at fasting baseline (BAS), 30m (30M) and 4h (4-H) in PREDICT. Normalised expression levels are INT(TPMs) residualised for potential confounders.

**Figure S11.** Expression signals replicated in the combined male and female sample from GOTO pre-intervention (**A**) and their expression trajectory in the PREDICT sample (**B**). In PREDICT changes were analysed at fasting baseline (BAS), 30m (30M) and 4h (4-H), and in GOTO changes were analysed at fasting baseline and 30m after meal (30M logFC). Directions of effect in PREDICT matched to GOTO at 30m (*) and 4h (^) are shown next to gene names. Normalised expression levels in PREDICT are INT(TPMs) residualised for potential confounders.

**Figure S12.** Diagram of overlapping genes with nominal signals in the epigenetic and expression postprandial results (*p* < 0.05). The overlap of epigenetic and expression trajectories (0-30m-4h) and the overlap of 4h epigenetic meta-analysis and 4h expression signals are represented in **A** and **B**, respectively. The peak 60 overlapping genes are described and ordered by the P-value of their epigenetic signal. The full list of overlapping genes is available in **Table S17**.

**Figure S13.** Nominal significance of eQTMs within target genes (*p* < 0.05). CpGs are ordered by position in the x-axis.

**Figure S14.** Co-methylation levels of probes found within target genes and correlation to gene expression. Blue and red indicate positive and negative correlation between probes while green and yellow indicate positive and negative correlation of DNAm probes to gene expression, respectively. Correlations were measured on DNAm and expression residuals using the Pearson’s method. The methylated position with the best correlation to gene expression levels is annotated in the figure. Genes that have nominally significant eQTMs in their loci (**Figure S13**) and best correlated probes that are eQTMs are highlighted in red.

**Figure S15.** Number of DNAm and expression signals previously reported under genetic regulation. Cis-meQTLs (*A*) for the main epigenetic signals of this study were identified in the MeQTL EPIC Database. Cis (*B*) and trans-eQTLs (*C*) for the main expression signals of this study were identified in the eQTLGen phase I database. Cis-eQTMs (*D*) for expression were identified in the BIOS database. All signals were selected with FDR = 5%.

**Figure S16.** Colocalization of genetic effects on the DNAm levels of cg00118229 and expression of *PDE9A*. **A**. meQTL and eQTL results from the MeQTL EPIC Database and eQTLGen phase I for the *PDE9A* locus, respectively. Target SNPs for were selected within 1 Mb of cg00118229 and the colocalised SNP rs2839581 (chr21:44,158,405) is highlighted. **B**. Postprandial trajectory of cg00118229 and associated phenotype result (*p* < 0.05).

**Figure S17.** Posterior probabilities for genetic colocalization sensitivity analysis. Posterior probabilities are shown as a function of *p*_12_ for cg00118229–PDE9A (**A**) and cg14880584–GPT2 (**B**). The green shaded region indicates the range of *p*_12_ values that support evidence of colocalization with (P(H4) > 0.8). Probabilities of H1, H2, H3 and H4 being true are represented in blue, green, light green and yellow dots, respectively.

**Table S1.** Meta-analysis of longitudinal DNAm changes associated with postprandial lipemia at 4h in PREDICT females and CORDIOPREV males (n = 225 participants; FDR = 5%).

**Table S2.** Enrichment analysis of postprandial lipemia DNAm meta-analysis results (FDR = 5%).

**Table S3.** EWAS Catalog trait enrichment of postprandial lipemia DNAm meta-analysis results.

**Table S4.** CD4+ T cell DNAm changes in the 27 DMPs associated with postprandial lipemia at 4h. Analysis was performed in a subset of PREDICT participants with CD4+ T cells collected postprandially (n = 10).

**Table S5.** Trajectory of signals associated with postprandial lipemia at 4h in PREDICT females (n = 104 participants).

**Table S6.** Epigenome-wide trajectory of DNAm changes associated with the postprandial state in PREDICT females (n = 104 participants; FDR = 5%).

**Table S7.** Sensitivity for menopause and hormone replacement therapy in analysis of DNAm changes associated with the postprandial state in PREDICT (n = 37 participants).

**Table S8.** Circadian oscillations of CpGs for the epigenetic signals of this study as reported by Oh et al. (2019).

**Table S9.** Enrichment analysis of trajectory of DNAm changes in PREDICT females (FDR = 5%).

**Table S10.** EWAS Catalog trait enrichment of trajectory of DNAm changes in PREDICT females.

**Table S11.** Replication of DNAm signals from Rask-Andersen et al. (2016) in PREDICT and CORDIOPREV postprandial DNAm analyses.

**Table S12.** Trajectory of gene expression changes associated with the postprandial state in PREDICT females (n = 50 participants; FDR = 5%).

**Table S13.** Sensitivity for analysis of gene expression changes associated with the postprandial state in PREDICT (n = 50 participants). Results are presented are from a linear mixed effects model adjusted for age, BMI, smoking, estimated blood cell proportions, season, RIN and GC mean as fixed effects, and rack, the participant, zygosity and family as random effects.

**Table S14.** Enrichment analysis of gene expression changes (FDR = 5%).

**Table S15.** Replication of gene expression signals from our analysis in the GOTO lifestyle intervention study.

**Table S16.** Gene expression signals from Rask-Andersen et al. (2016) in the PREDICT gene expression trajectory analysis.

**Table S17.** List of overlapped genes with nominal signals in the epigenetic and expression postprandial results (*p* < 0.05).

**Table S18.** Enrichment analysis of overlapped genes with nominal signals in the epigenetic and expression postprandial results (FDR = 25%).

**Table S19.** Cis-meQTLs identified in the EPIC MeQTL Database for the epigenetic signals of this study (FDR = 5%).

**Table S20.** Cis and trans-eQTLs identified in the eQTLGen phase I database for the main expression trajectory signals of this study (FDR = 5%).

**Table S21.** Cis-eQTMs identified in the BIOS database for the main expression trajectory signals of this study (FDR = 5%).

**Table S22.** GWAS Catalog traits identified for the main epigenetic and expression signals of this study.

**Table S23.** DNAm signals associated with plasma glucose and triglycerides postprandially in PREDICT females (n_max_ = 104 participants; FDR = 5%).

**Table S24.** Subset of gene expression signals associated with plasma glucose and triglycerides postprandially in PREDICT females (n_max_ = 50 participants; FDR = 5%). Results presented are from a subset of genes where DNAm changes with plasma glucose and triglycerides were also observed in the same set of participants.

**Table S25.** Replication of DNAm signals from Lai et al. (2016) in PREDICT analyses of postprandial triglycerides.

**Table S26.** Epigenetic aging markers associated with plasma glucose and triglycerides postprandially in PREDICT females (n_max_ = 104 participants).

**Table S27.** Gene expression results for DNAm signals from Lai et al. (2016) in PREDICT analyses of postprandial triglycerides.

**Table S28.** Postprandial glucose and triglyceride levels in the PREDICT sample.

