## Supplemental Text and Figures for "DNA methylation and gene expression trajectories of human postprandial metabolism"

### Supplementary Information

#### Notes on “Longitudinal meta-analysis identifies DNA methylation signals postprandially at 4 hours”

The results are presented after multiple testing correction (FDR 5%) and accounting for heterogeneity (HetISq < 85% and HetPval ≥ 0.01). We reported 108 DMPs from our main meta-analysis. Five additional sites were differentially methylated at 4h, but these results did not surpass meta-analysis heterogeneity filters. Nonetheless, we report them in Table S1, but where HetPASS column equals “No”.

Sensitivity analysis of results was performed using DNAm β-values. Unlike our main DNAm M-value analysis, peak signals identified with raw DNAm β-values suffered from high heterogeneity (*i.e.:* only 12 of the 50 (24%) peak signals passed meta-analysis heterogeneity filters).

#### Notes on “Genetic colocalization analysis”

Bayesian colocalization analysis quantifies the probability of a given SNP being causally associated with two traits. This method requires prior probabilities for a SNP being associated with trait 1 only (*p_1_*), trait 2 only (*p_2_*), and both traits (*p_12_*)^1–3^. We specified these probabilities for each CpG–gene comparison independently. We approximated priors based on the number of SNPs with FDR < 5% in each locus relative to the total number of tests (representing *q_1_ = p_1_ + p_12_* and *q_2_ = p_2_ + p_12_* for each trait, respectively). Additionally, priors were estimated from the Pearson correlation of effect sizes across LD-clumped SNPs (*p_12_ = |r|* min(*q_1_, q_2_*)). If either *p_1_, p_2_* or *p_12_* exceeded 1/(n_SNP_+1) (where n_SNP_ is the number of SNPs available for both traits in the locus), prior probabilities were scaled to this maximum value. The table below displays the number of available SNPs and specified priors for colocalization analysis. Phenotypic variances for each trait are required for colocalization. These were estimated with a modified version of the R function *estimate_trait_sd* from the *TwoSampleMR* R package^4^.

**Table.** CpG-gene pairs, number of SNPs and values of priors estimated for colocalization analysis of postprandial DNAm and gene expression.

| **CpG** | **Gene** | ***n*_SNP_^a^** | ***p*_1_ (CpG)** | ***p*_2_ (Gene)** | ***p*_12_** |
| --- | --- | --- | --- | --- | --- |
| cg00118229 | *PDE9A* | 5279 | 1.5E-06 | 1.9E-04 | 2.0E-06 |
| cg00525681 | *SLITRK5* | 4126 | 7.6E-05 | 2.4E-04 | 1.1E-04 |
| cg04340435 | *TMCO3* | 3627 | 1.9E-05 | 2.8E-04 | 1.2E-05 |
| cg04340435 | *DCUN1D2* | 3656 | 9.8E-05 | 2.7E-04 | 8.4E-05 |
| cg04951739 | *ZNF70* | 3005 | 3.3E-04 | 2.0E-04 | 1.9E-05 |
| cg07235635 | *GATAD2A* | 3000 | 5.2E-06 | 3.3E-04 | 1.9E-05 |
| cg08111040 | *C9orf89* | 4157 | 2.1E-05 | 2.4E-04 | 1.7E-05 |
| cg08939787 | *RAMP1* | 4762 | 5.3E-05 | 2.1E-04 | 8.3E-05 |
| cg10163682 | *ATP7B* | 2784 | 8.1E-05 | 3.6E-04 | 5.0E-06 |
| cg11139891 | *EFTUD2* | 3872 | 5.9E-05 | 2.6E-04 | 3.9E-05 |
| cg11410920 | *SYS1* | 3506 | 1.4E-05 | 2.9E-04 | 2.1E-05 |
| cg11996592 | *ZC3H3* | 3671 | 2.7E-05 | 2.7E-04 | 2.5E-05 |
| cg14880584 | *GPT2* | 253 | 8.4E-04 | 8.6E-04 | 2.5E-03 |
| cg17108629 | *RAB38* | 4599 | 1.9E-05 | 2.2E-04 | 2.4E-05 |
| cg17656260 | *ZNF790* | 2927 | 2.2E-04 | 1.5E-04 | 3.4E-04 |
| cg19616459 | *ABCC1* | 2350 | 5.8E-05 | 4.3E-04 | 6.5E-06 |
| cg20981127 | *NR2F6* | 3611 | 1.3E-05 | 2.8E-04 | 4.1E-05 |
| cg21150537 | *KCNG2* | 3242 | 8.2E-05 | 3.1E-04 | 2.3E-04 |
| cg26803385 | *CROCC* | 3351 | 4.7E-05 | 3.0E-04 | 2.1E-06 |

*^a^Number of SNPs available for both traits in the locus*

#### Notes on “Baseline DNA methylation signals of postprandial glucose and triglycerides trajectories”

In addition to using baseline EEA measures to predict the interindividual variation of glucose and triglycerides postprandially, we also estimated whether measures of epigenetic aging themselves changed immediately after meal. Some variation in EAA estimates were observed postprandially but these were not statistically significantly in association with the timepoints. Accordingly, we used only baseline estimates in our analysis.

#### Figures

(next)

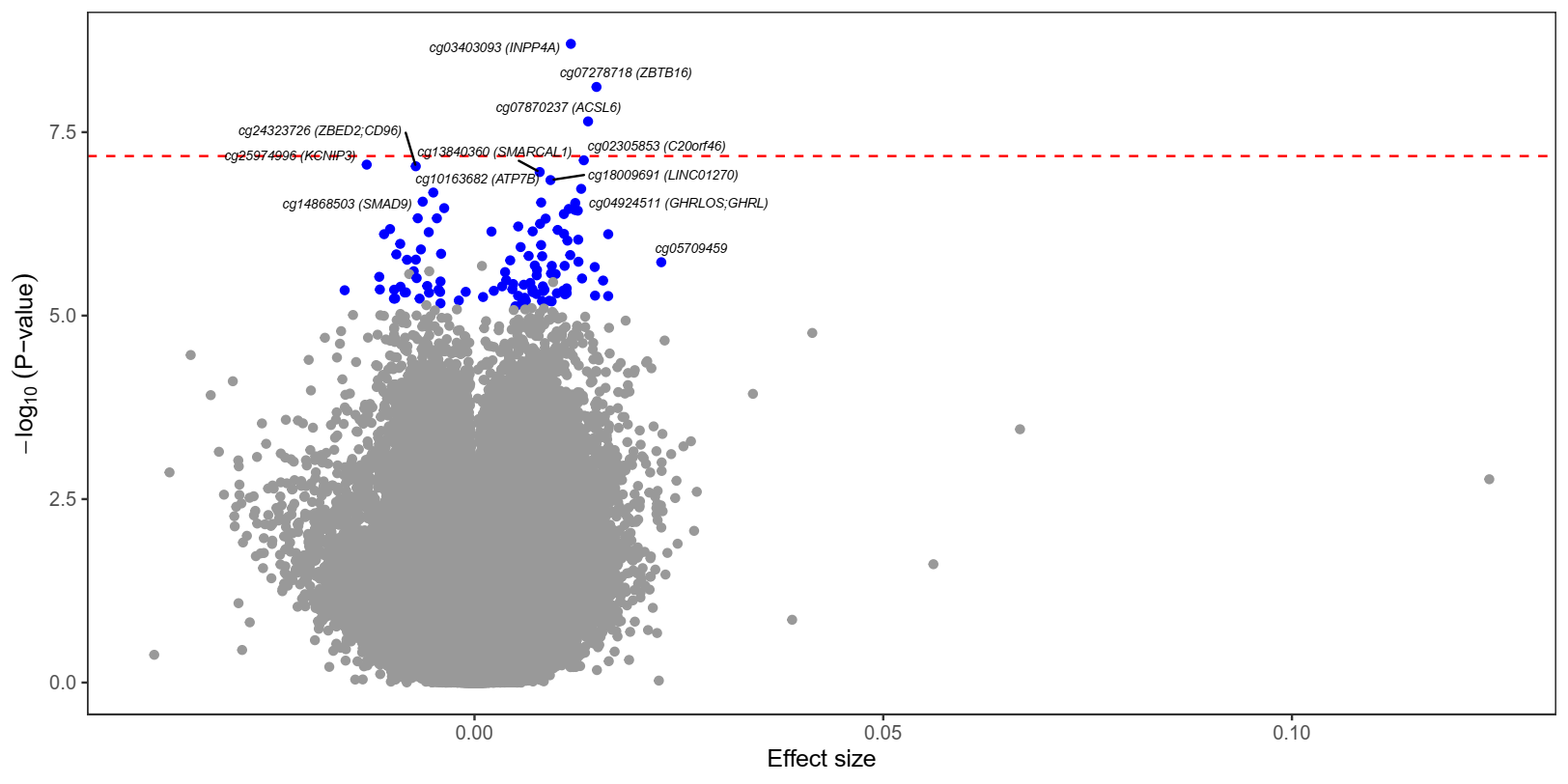

**Figure S1.** Alternative plot of DNAm signals identified at 4h postprandially in PREDICT and CORDIOPREV (n = 225 participants). The effect sizes displayed here are based on DNAm β-values instead of M-values (**Figure 2A** from the main text).

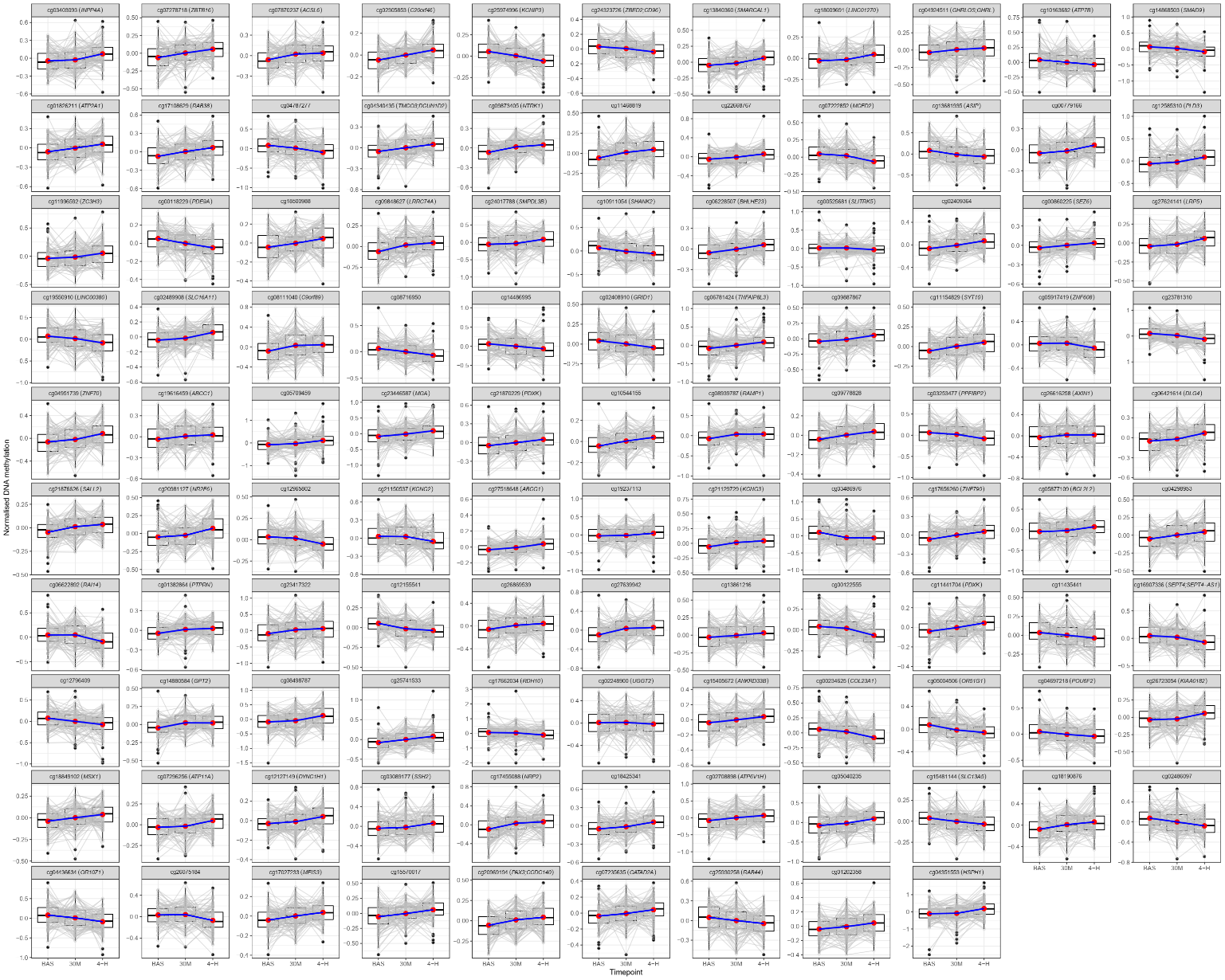

**Figure S2.** Postprandial DNAm trajectories of the 108 DMPs identified in the 4h epigenetic meta-analysis. Changes were analysed at fasting baseline (BAS), 30m (30M) and 4h (4-H) in PREDICT. Normalised DNAm levels are DNAm M-values residualised for potential confounders.

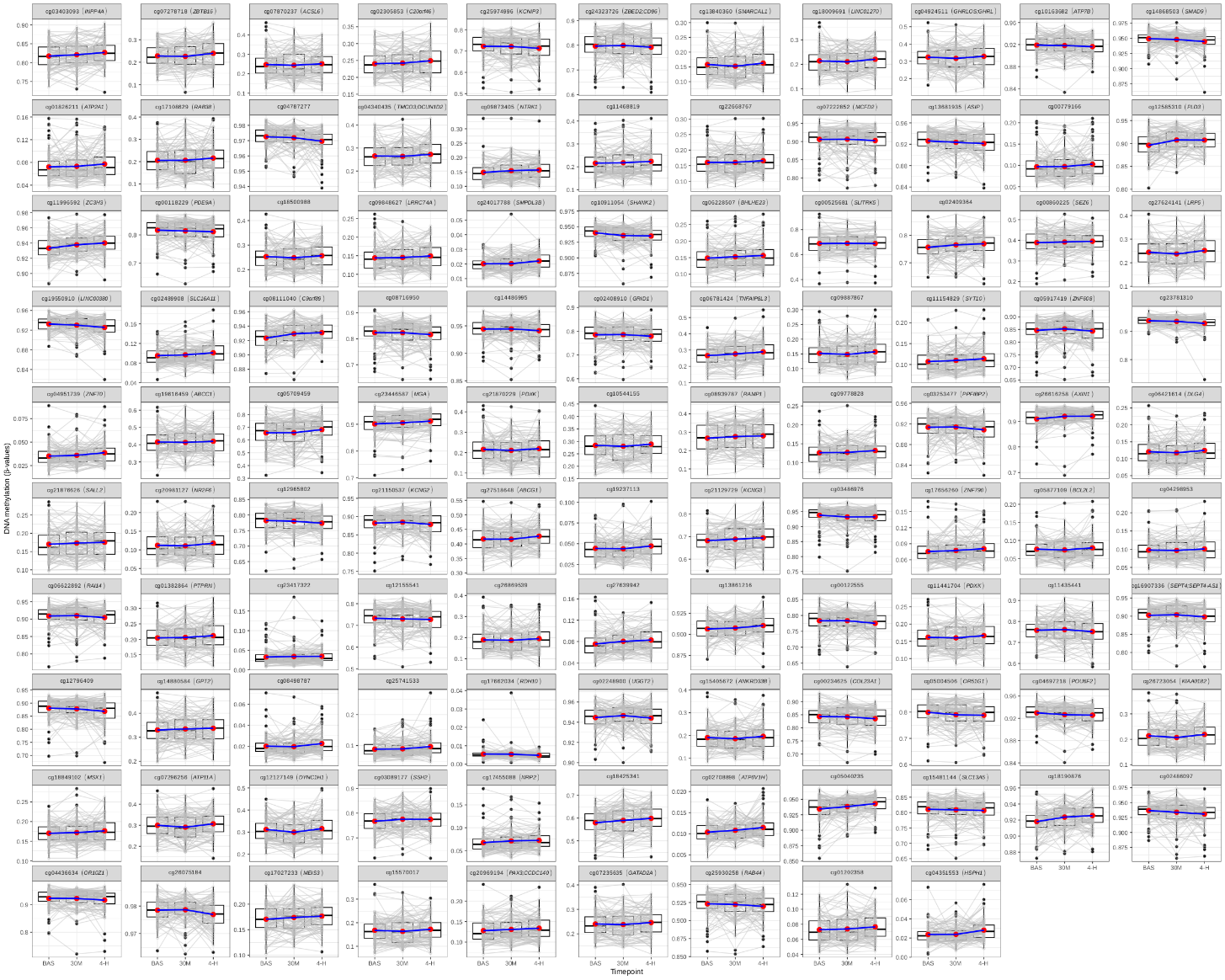

**Figure S3.** Postprandial DNAm trajectories of the 108 DMPs identified in the 4h epigenetic meta-analysis. Changes were analysed at fasting baseline (BAS), 30m (30M) and 4h (4-H) in PREDICT. DNAm levels are raw unadjusted DNAm β-values.

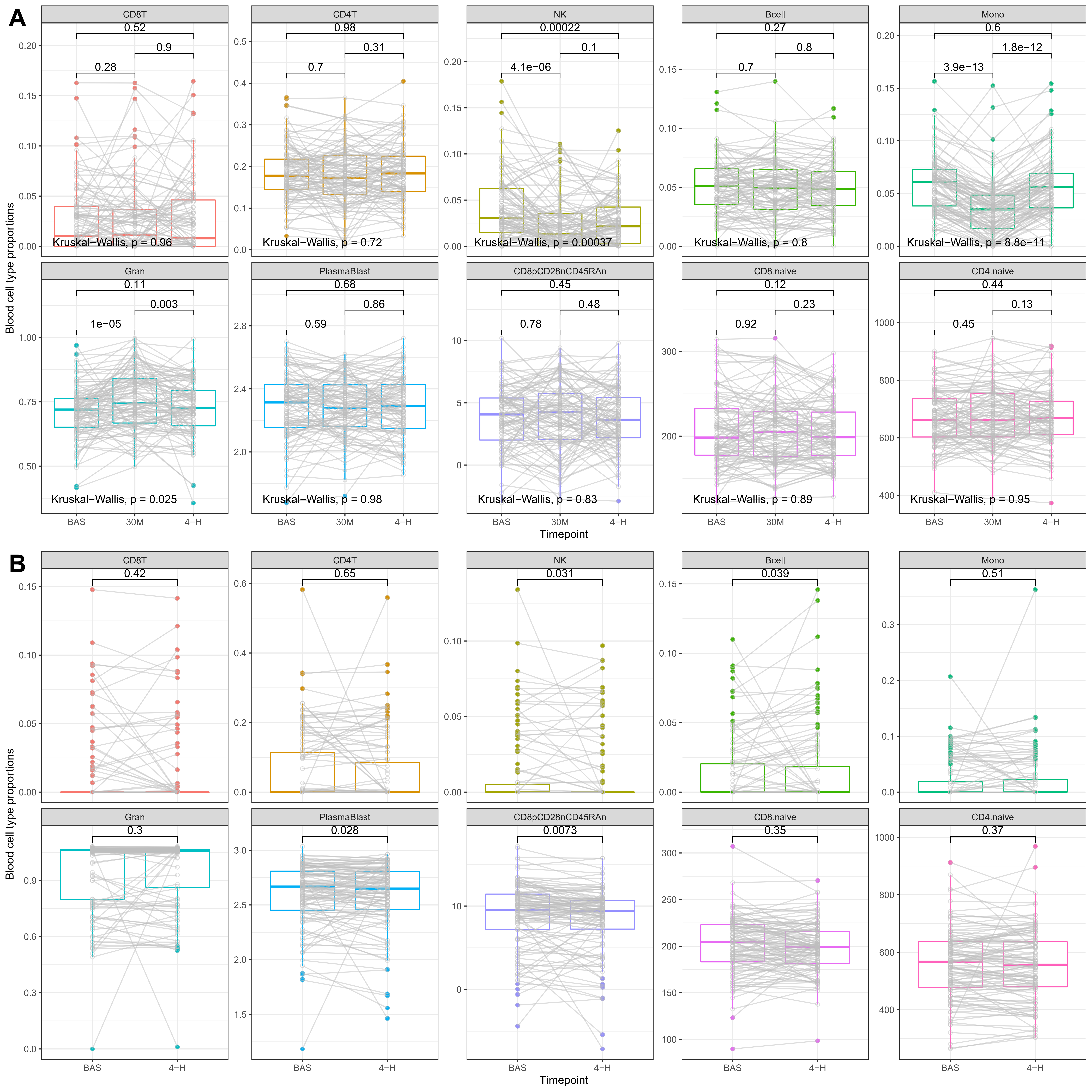

**Figure S4.** Postprandial blood cell proportion changes in the PREDICT (**A**) and COPRDIOPEV (**B**) studies. Changes were analysed at fasting baseline (BAS), 30m (30M) and 4h (4-H) after meal using the Wilcoxon signed rank test for each pair of samples and the Kruskal-Wallis test for overall differences in PREDICT. Estimated blood cell proportions are based on DNAm patterns from whole blood in PREDICT and polymorphonuclear leukocyte blood fractions in CORDIOPREV.

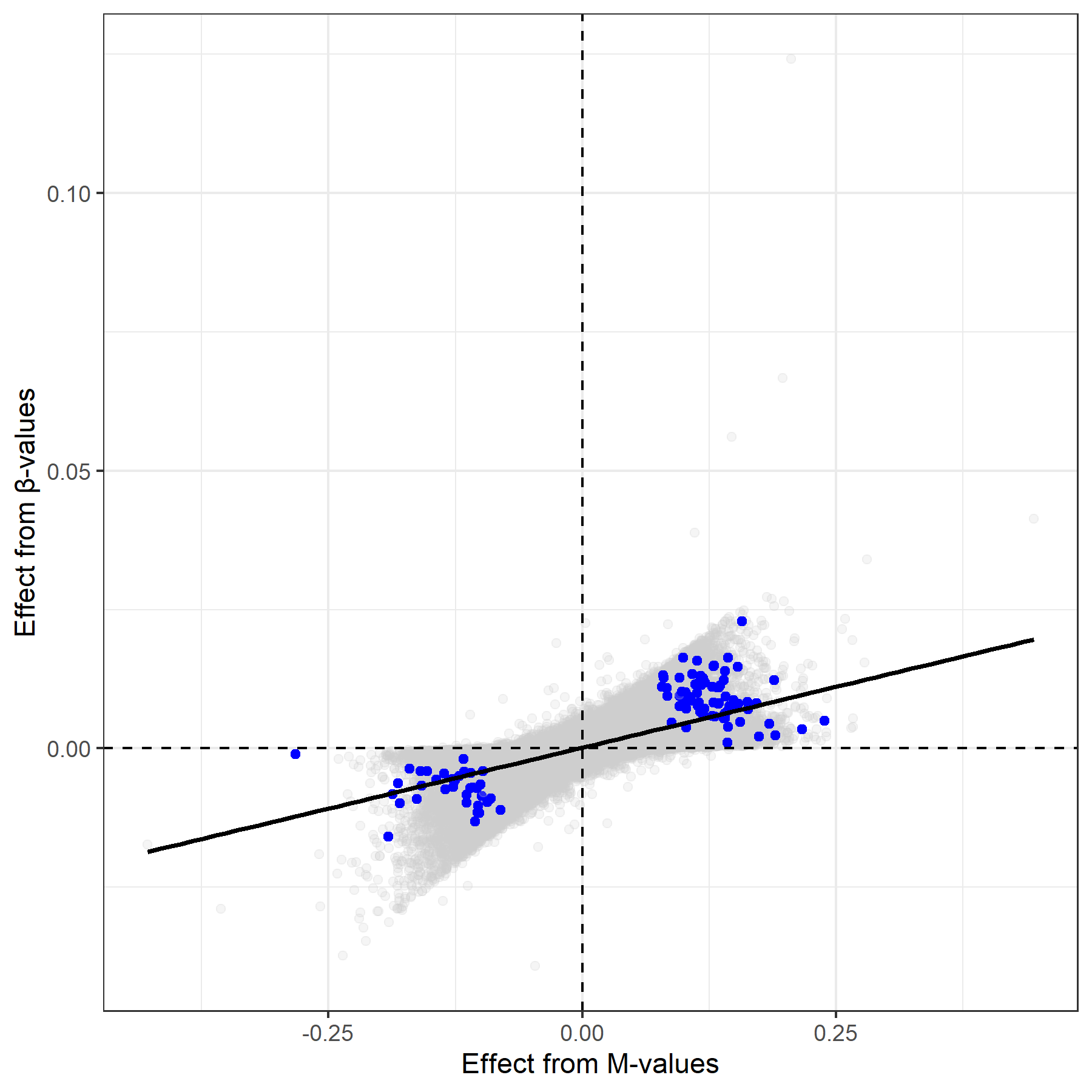

**Figure S5.** Directions of effect at 4h postprandially using M- or β-values as the response variable to timepoint in the PREDICT and CORDIOPREV cohorts (n = 225 participants). Signals from the main postprandial DNAm 4h meta-analysis (**Figure 2A; Table S1**) are highlighted in blue.

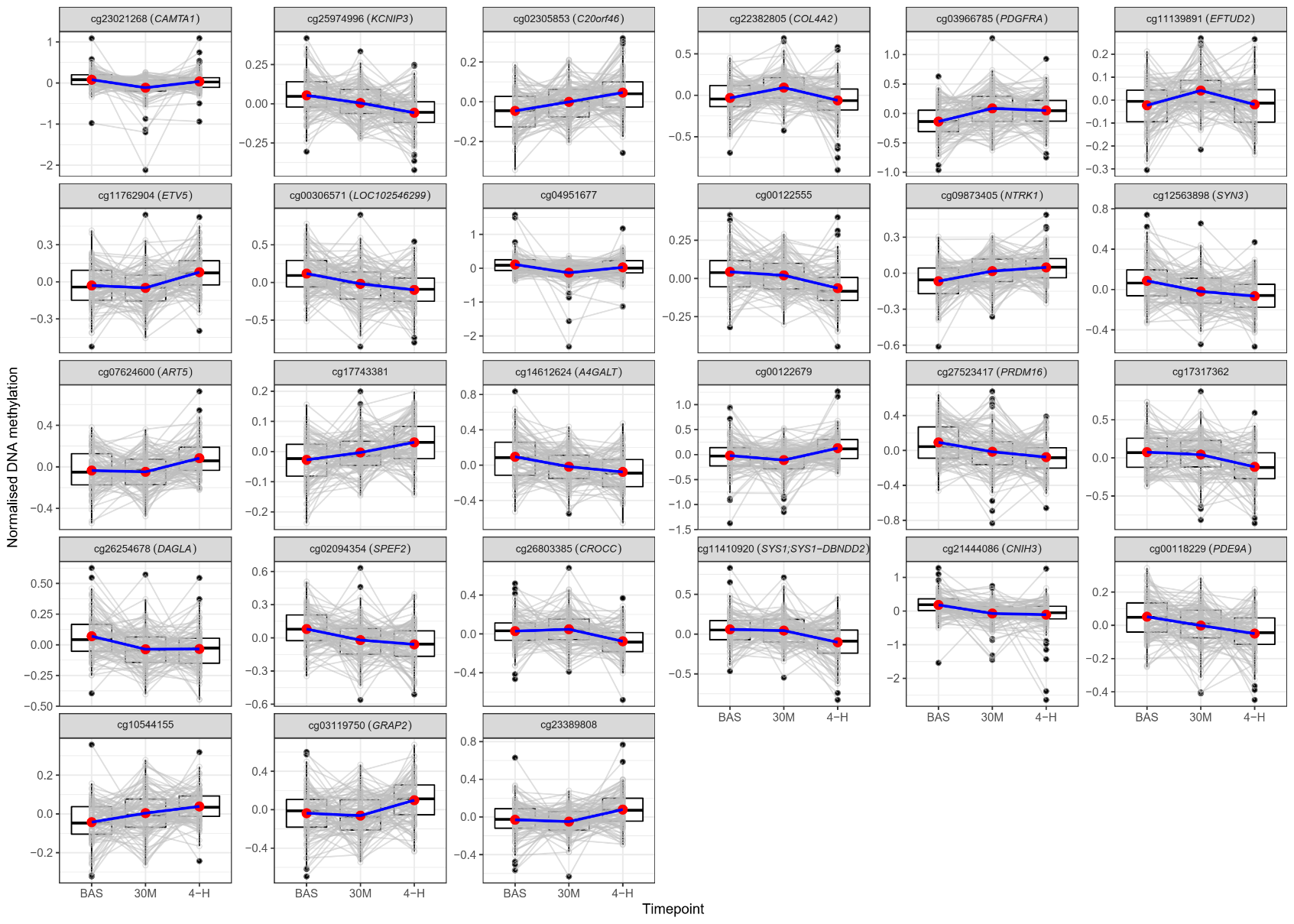

**Figure S6.** Postprandial DNAm trajectories of the 27 DMPs identified in the main epigenetic trajectory analysis. Changes were analysed at fasting baseline (BAS), 30m (30M) and 4h (4-H) in PREDICT. Normalised DNAm levels are DNAm M-values residualised for potential confounders.

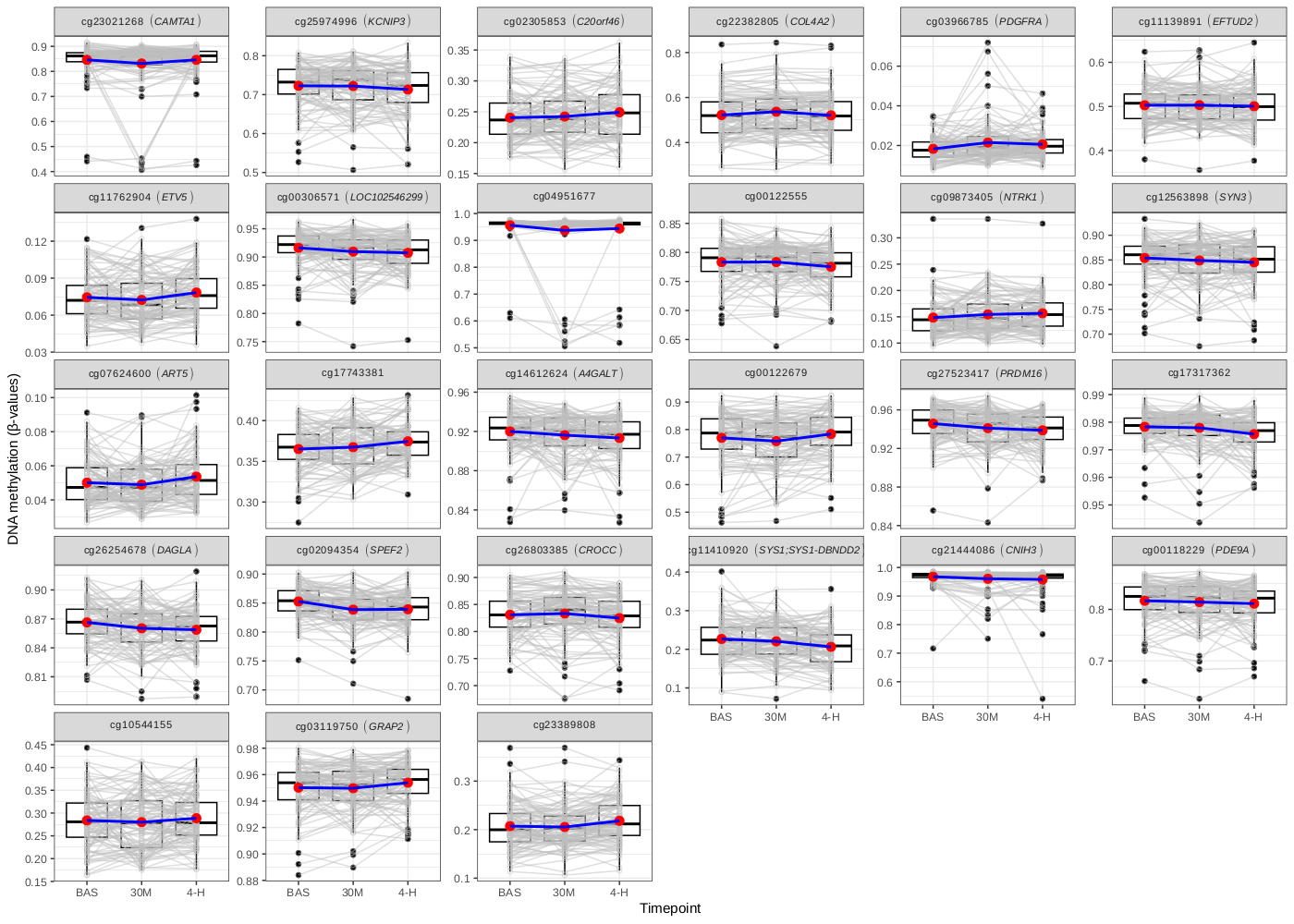

**Figure S7.** Postprandial DNAm trajectories of the 27 DMPs identified in the main epigenetic trajectory analysis. Changes were analysed at fasting baseline (BAS), 30m (30M) and 4h (4-H) in PREDICT. DNAm levels are raw unadjusted DNAm β-values.

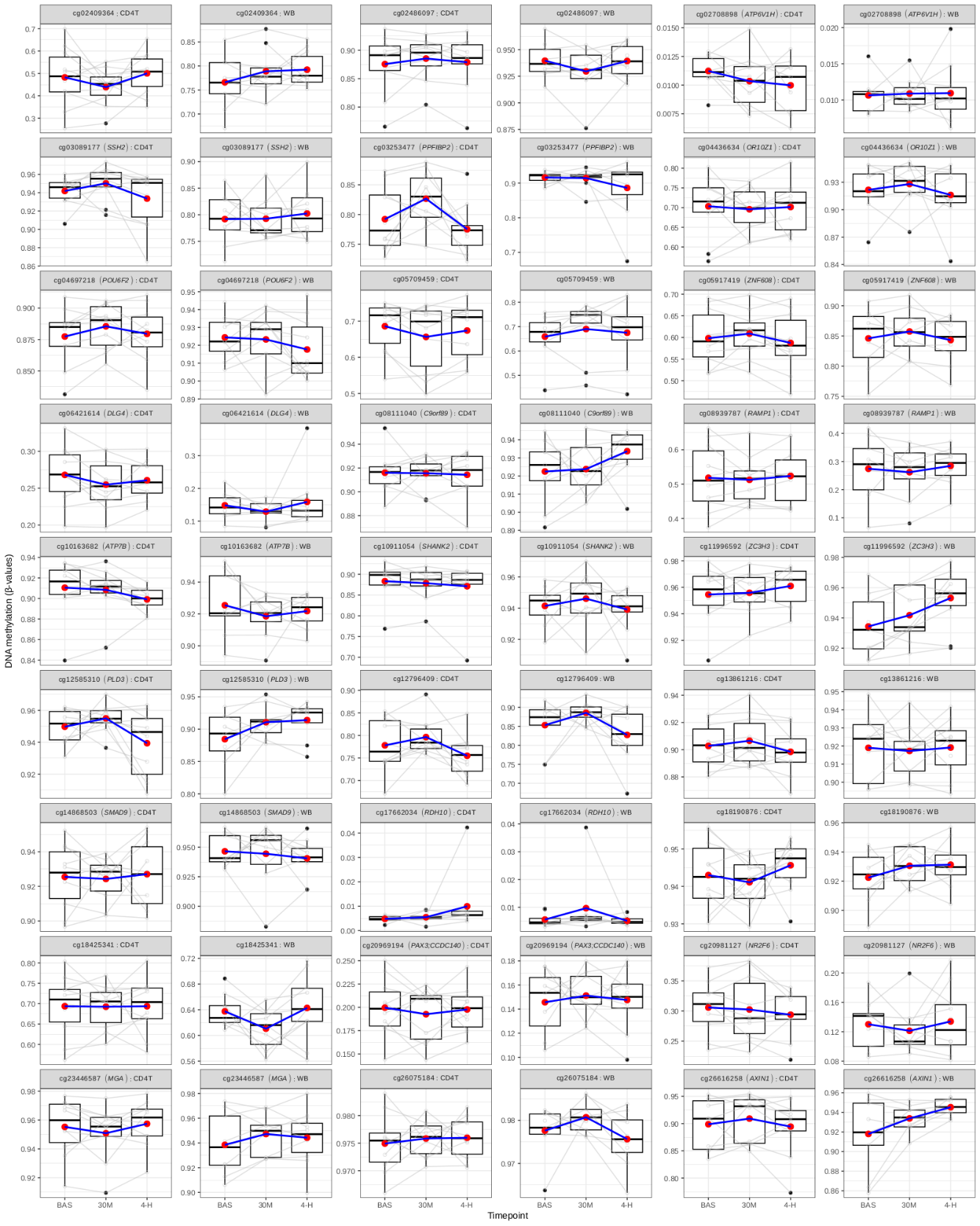
 **Figure S8.** Postprandial DNAm trajectories in CD4+ T cells (CD4T) and matched whole blood (WB) samples. Trajectories are represented for signals from the main postprandial lipemia meta-analysis (**Figure 1, Table S1**) with consistent directions of effect in the whole blood raw DNAm β-values from the 10-participant subset with CD4+ T cells available.

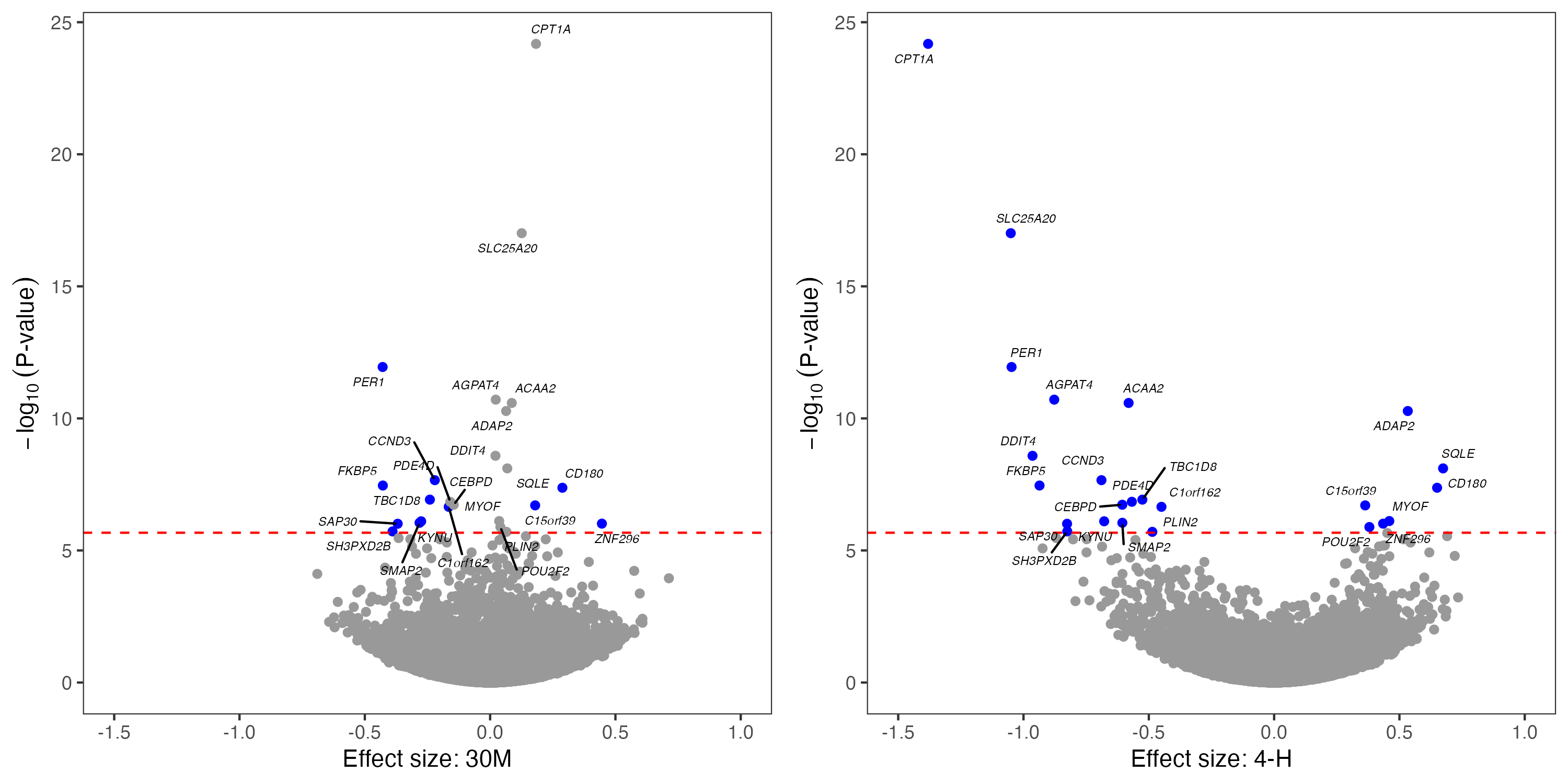

**Figure S9.** Volcano plots of gene expression signals identified at 30m (left) and 4h (right) postprandially in PREDICT (n = 50 participants).

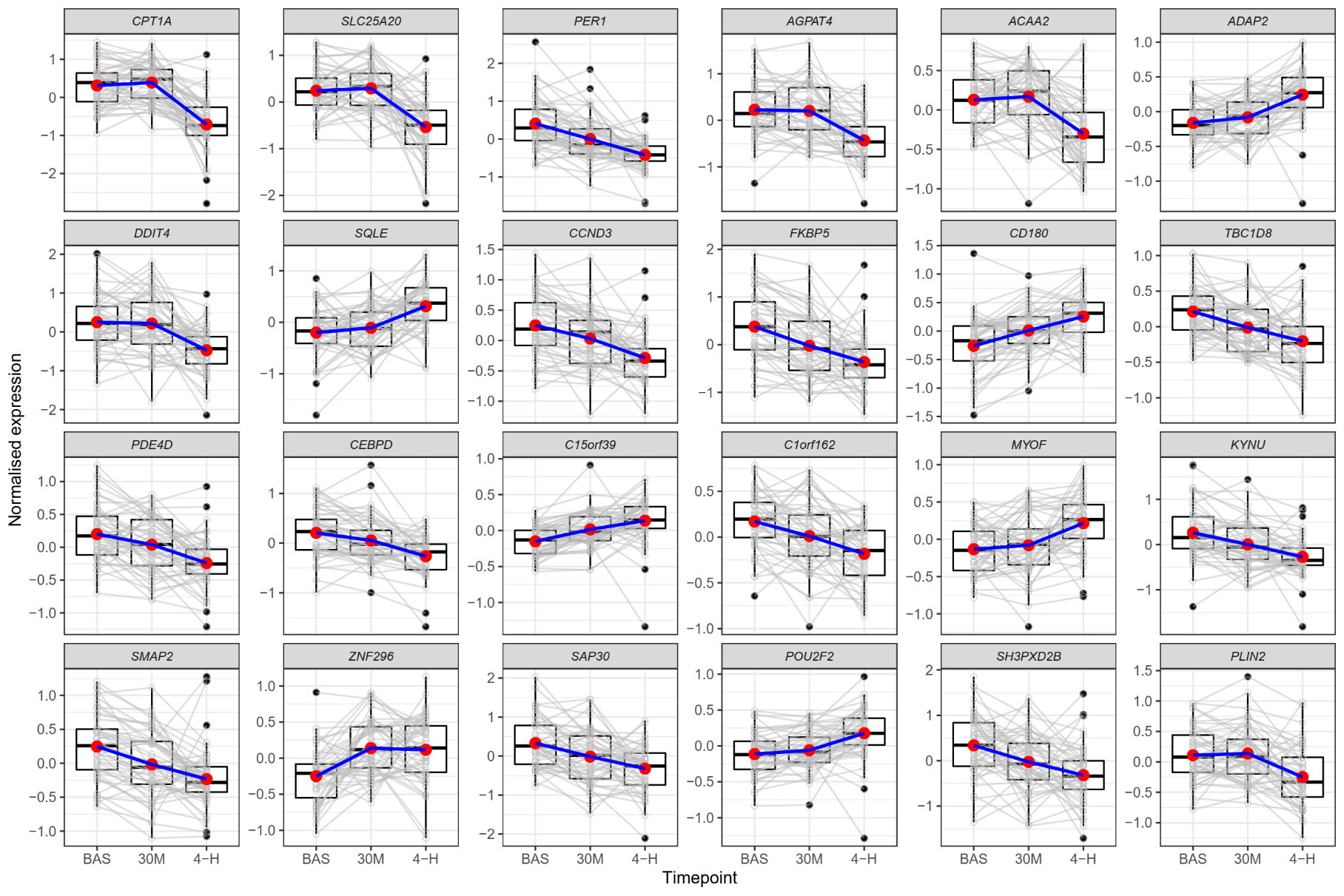

**Figure S10.** Postprandial expression trajectories of the 24 genes identified in the main expression trajectory analysis. Changes were analysed at fasting baseline (BAS), 30m (30M) and 4h (4-H) in PREDICT. Normalised expression levels are INT(TPMs) residualised for potential confounders.

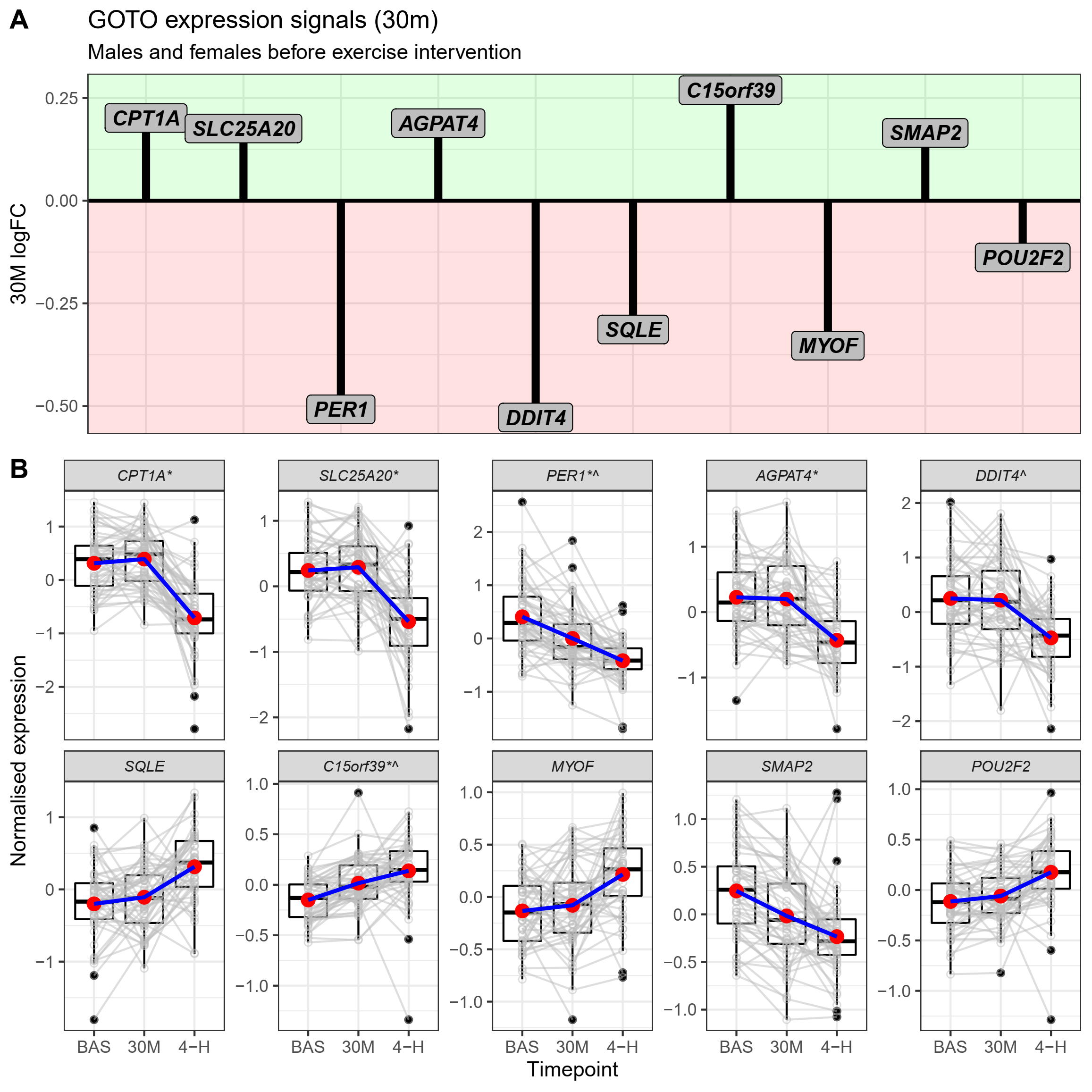

**Figure S11.** Expression signals replicated in the combined male and female sample from GOTO pre-intervention (**A**) and their expression trajectory in the PREDICT sample (**B**). In PREDICT changes were analysed at fasting baseline (BAS), 30m (30M) and 4h (4-H), and in GOTO changes were analysed at fasting baseline and 30m after meal (30M logFC). Directions of effect in PREDICT matched to GOTO at 30m (*) and 4h (^) are shown next to gene names. Normalised expression levels in PREDICT are INT(TPMs) residualised for potential confounders.

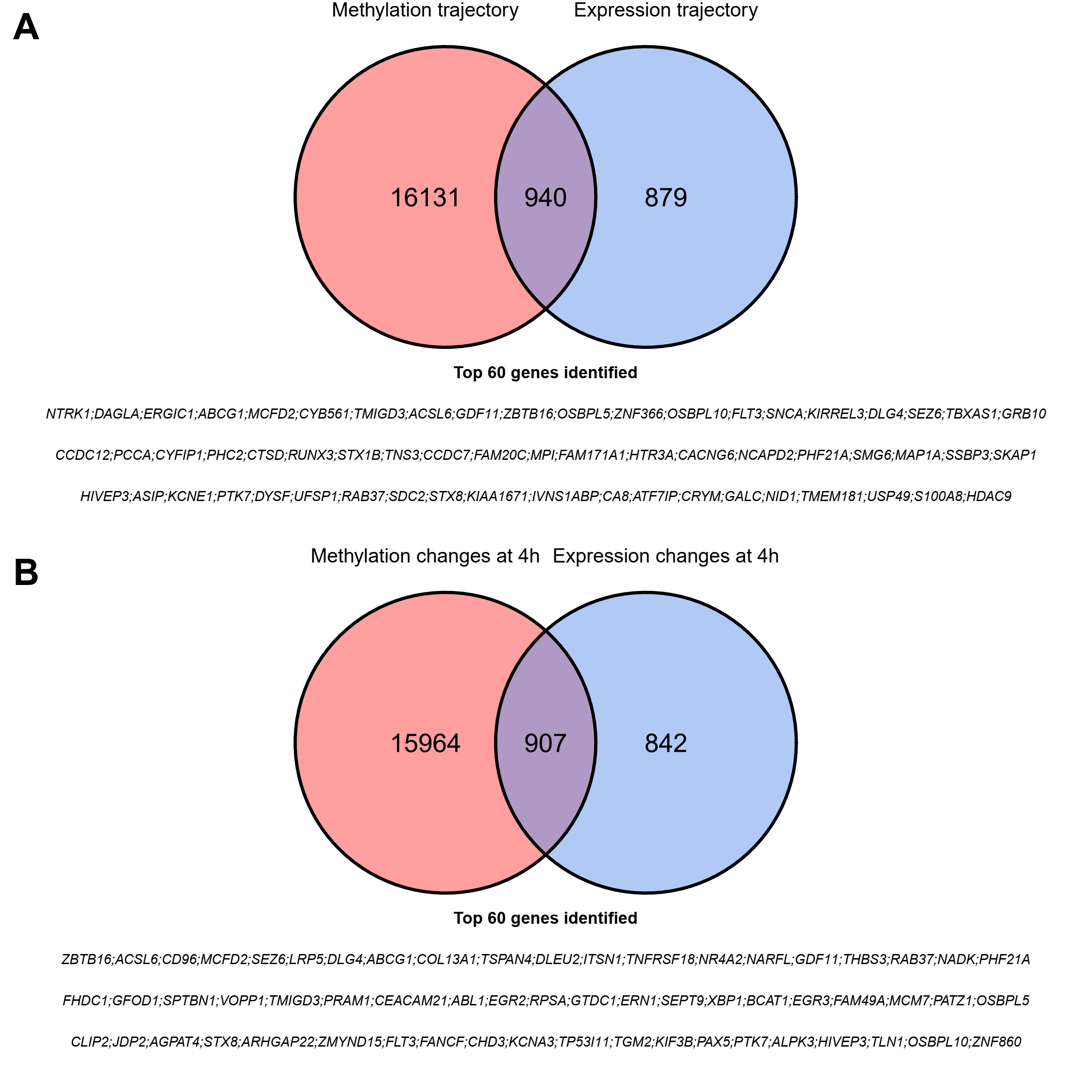

**Figure S12.** Diagram of overlapping genes with nominal signals in the epigenetic and expression postprandial results (*p* < 0.05). The overlap of epigenetic and expression trajectories (0-30m-4h) and the overlap of 4h epigenetic meta-analysis and 4h expression signals are represented in **A** and **B**, respectively. The peak 60 overlapping genes are described and ordered by the P-value of their epigenetic signal. The full list of overlapping genes is available in **Table S17.**

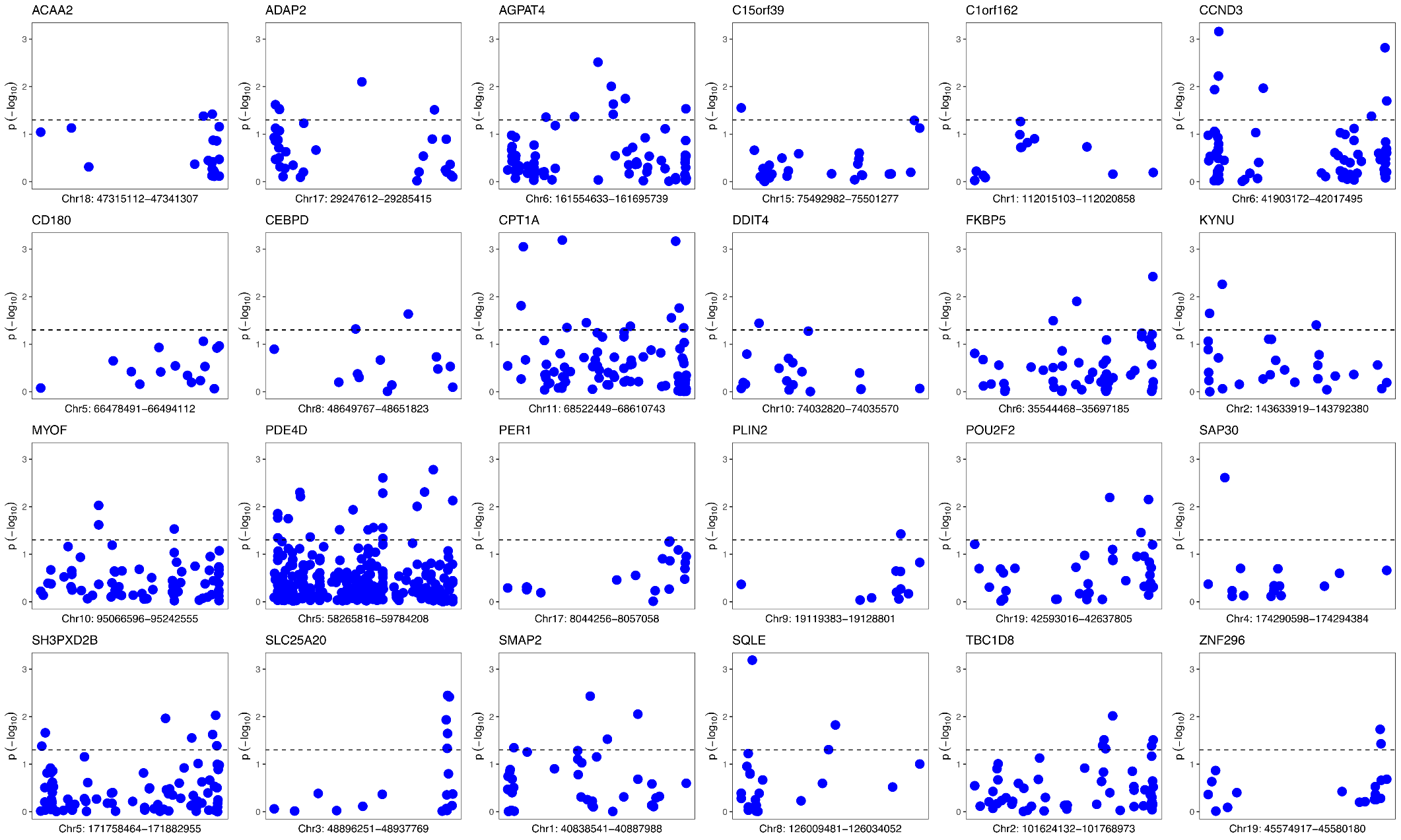

**Figure S13.** Nominal significance of eQTMs within target genes (*p* < 0.05). CpGs are ordered by position in the x-axis.

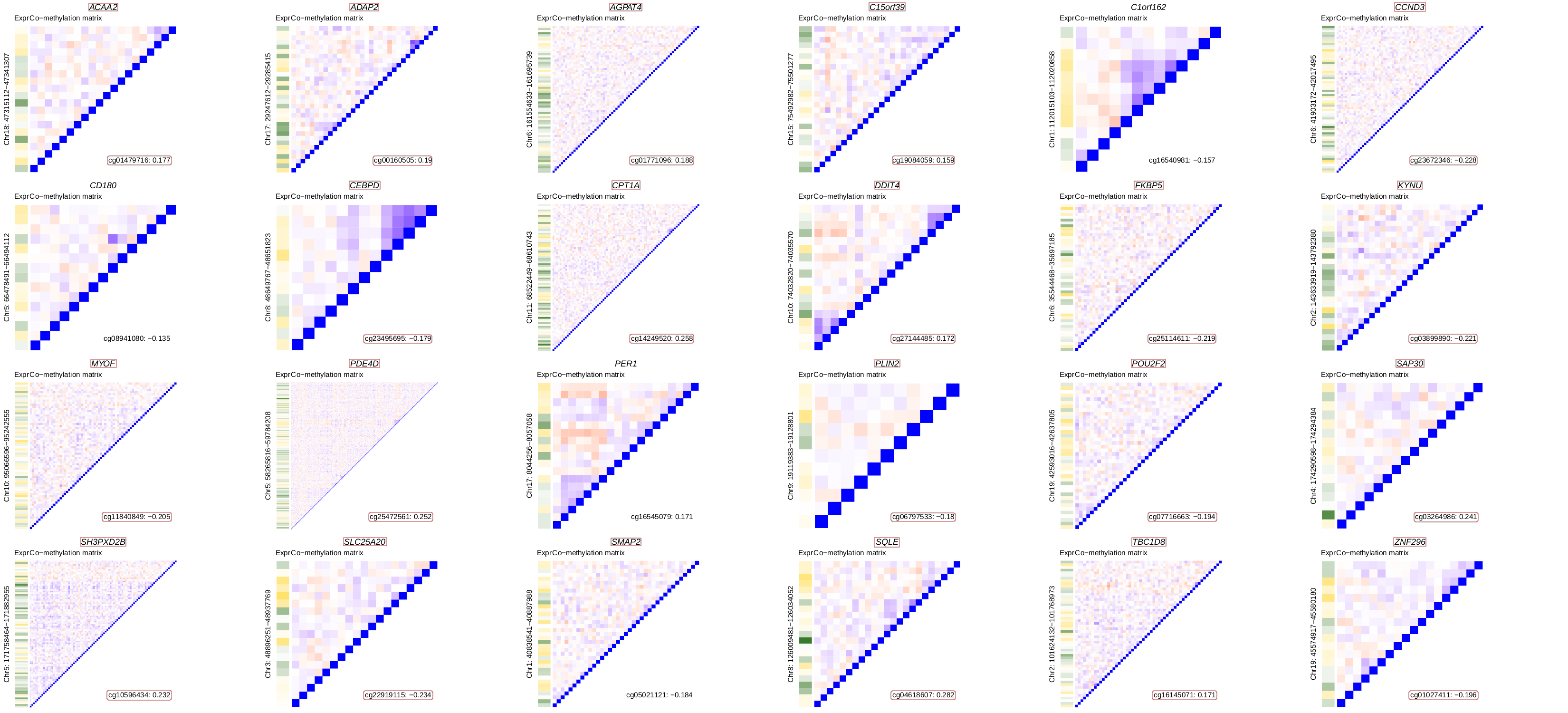

**Figure S14.** Co-methylation levels of probes found within target genes and correlation to gene expression. Blue and red indicate positive and negative correlation between probes while green and yellow indicate positive and negative correlation of DNAm probes to gene expression, respectively. Correlations were measured on DNAm and expression residuals using the Pearson’s method. The methylated position with the best correlation to gene expression levels is annotated in the figure. Genes that have nominally significant eQTMs in their loci (**Figure S13**) and best correlated probes that are eQTMs are highlighted in red.

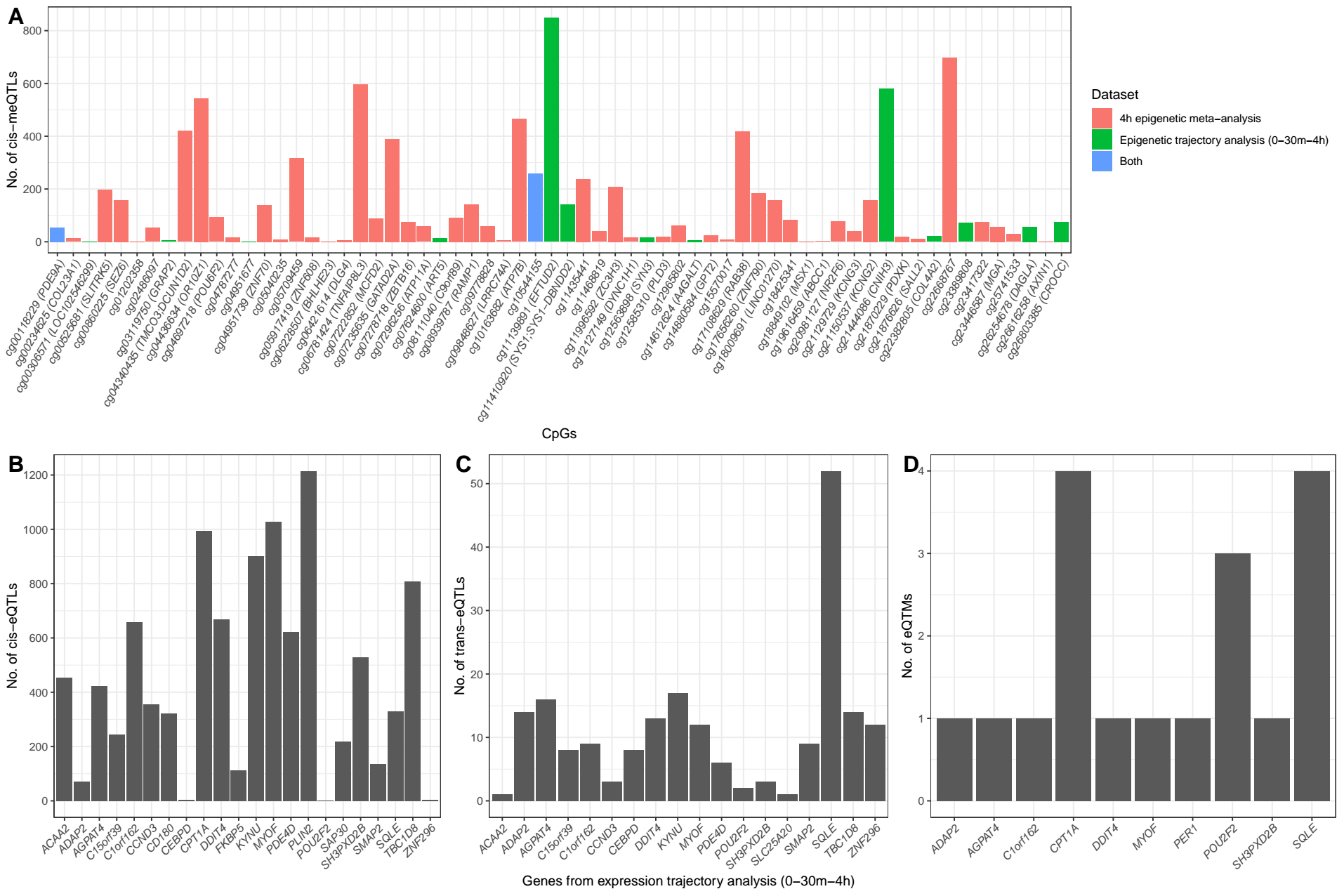

**Figure S15.** Number of DNAm and expression signals previously reported under genetic regulation. Cis-meQTLs (**A**) for the main epigenetic signals of this study were identified in the MeQTL EPIC Database. Cis (**B**) and trans-eQTLs (**C**) for the main expression signals of this study were identified in the eQTLGen phase I database. Cis-eQTMs (**D**) for expression were identified in the BIOS database. All signals were selected with FDR = 5%.

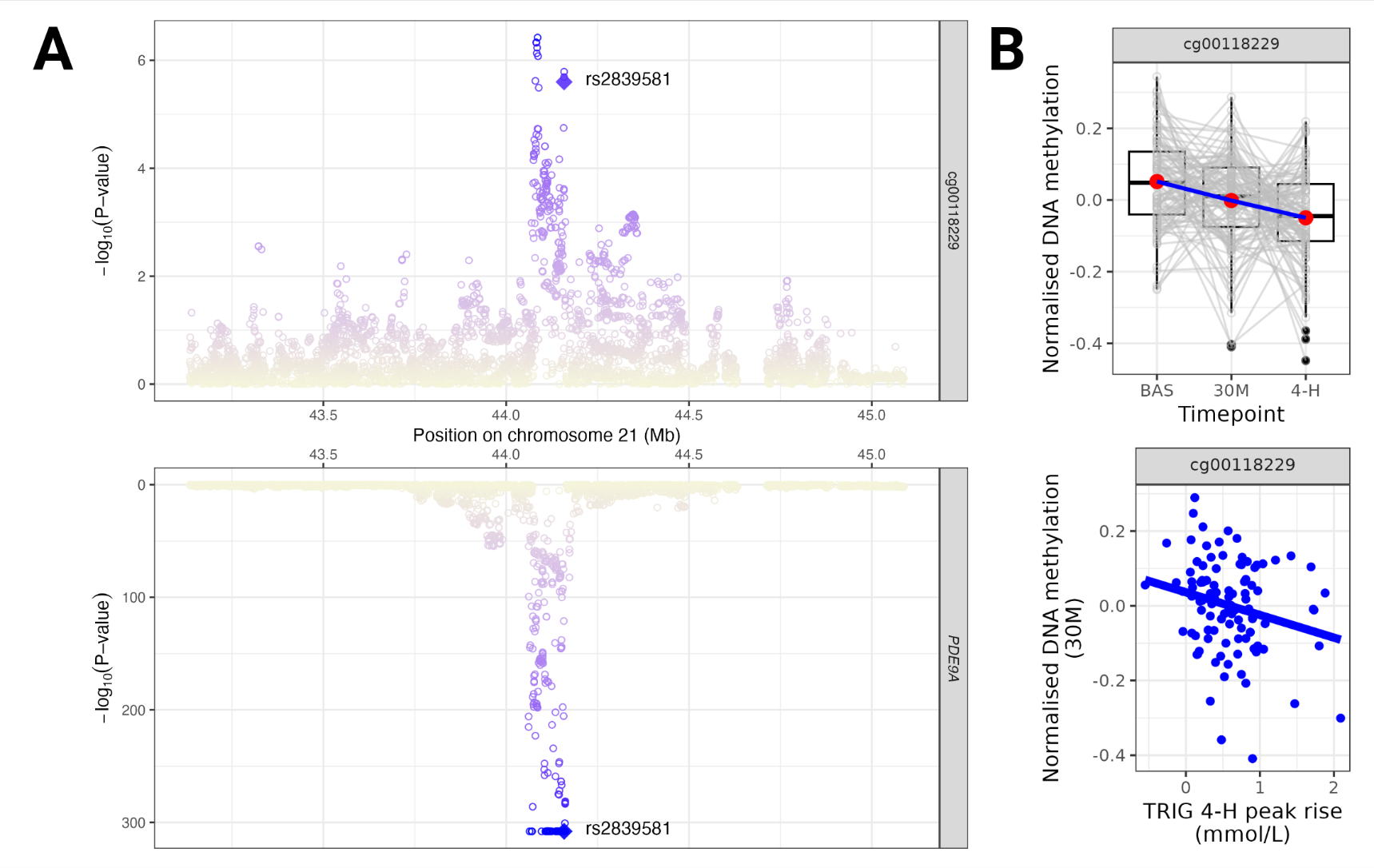

**Figure S16.** Colocalization of genetic effects on the DNAm levels of cg00118229 and expression of *PDE9A*. **A.** meQTL and eQTL results from the MeQTL EPIC Database and eQTLGen phase I for the *PDE9A* locus, respectively. Target SNPs for were selected within 1 Mb of cg00118229 and the colocalised SNP rs2839581 (chr21:44,158,405) is highlighted. **B.** Postprandial trajectory of cg00118229 and associated phenotype result (*p* < 0.05).

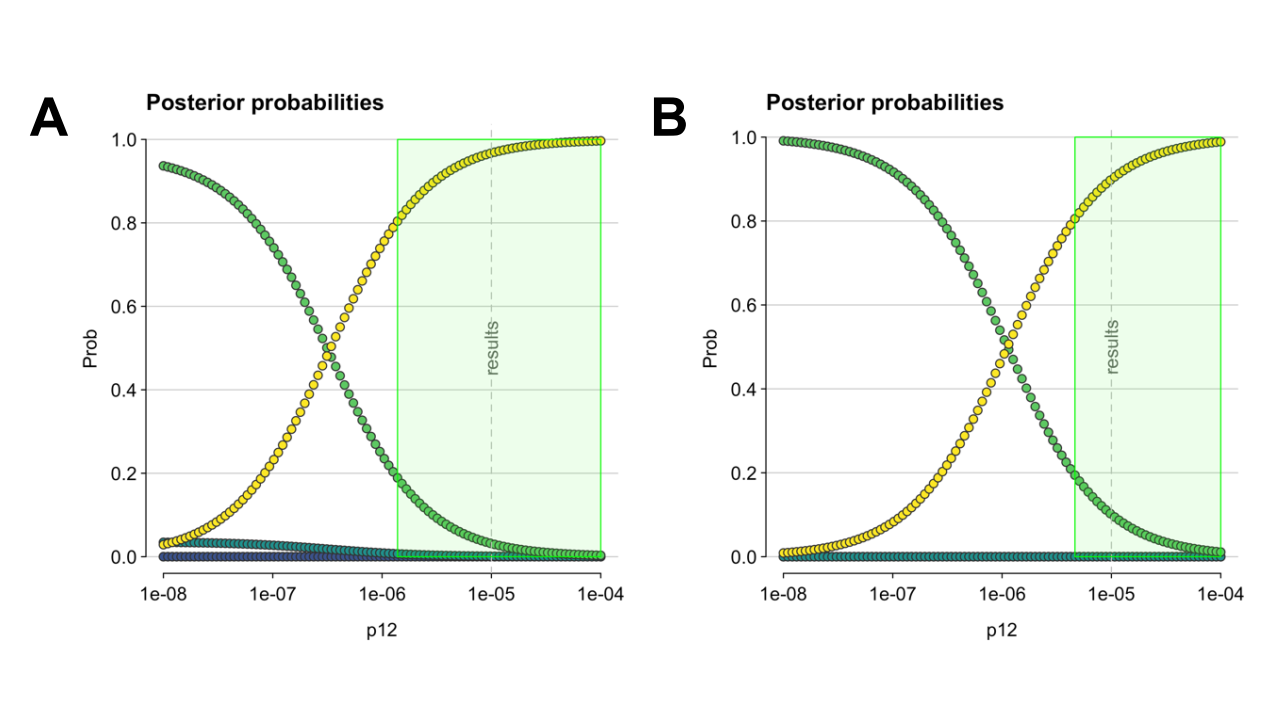

**Figure S17.** Posterior probabilities for genetic colocalization sensitivity analysis. Posterior probabilities are shown as a function of *p*_12_ for cg00118229–*PDE9A* (**A**) and cg14880584–*GPT2* (**B**). The green shaded region indicates the range of *p_12_* values that support evidence of colocalization with (P(H4) > 0.8). Probabilities of H1, H2, H3 and H4 being true are represented in blue, green, light green and yellow dots, respectively.

.
